# Dual Proteome-scale Networks Reveal Cell-specific Remodeling of the Human Interactome

**DOI:** 10.1101/2020.01.19.905109

**Authors:** Edward L. Huttlin, Raphael J. Bruckner, Jose Navarrete-Perea, Joe R. Cannon, Kurt Baltier, Fana Gebreab, Melanie P. Gygi, Alexandra Thornock, Gabriela Zarraga, Stanley Tam, John Szpyt, Alexandra Panov, Hannah Parzen, Sipei Fu, Arvene Golbazi, Eila Maenpaa, Keegan Stricker, Sanjukta Guha Thakurta, Ramin Rad, Joshua Pan, David P. Nusinow, Joao A. Paulo, Devin K. Schweppe, Laura Pontano Vaites, J. Wade Harper, Steven P. Gygi

**Author notes:** Correspondence (E.L.H.), (J.W.H.), (S.P.G.).

## Abstract

Thousands of interactions assemble proteins into modules that impart spatial and functional organization to the cellular proteome. Through affinity-purification mass spectrometry, we have created two proteome-scale, cell-line-specific interaction networks. The first, BioPlex 3.0, results from affinity purification of 10,128 human proteins – half the proteome – in 293T cells and includes 118,162 interactions among 14,586 proteins; the second results from 5,522 immunoprecipitations in HCT116 cells. These networks model the interactome at unprecedented scale, encoding protein function, localization, and complex membership. Their comparison validates thousands of interactions and reveals extensive customization of each network. While shared interactions reside in core complexes and involve essential proteins, cell-specific interactions bridge conserved complexes, likely ‘rewiring’ each cell’s interactome. Interactions are gained and lost in tandem among proteins of shared function as the proteome remodels to produce each cell’s phenotype. Viewable interactively online through BioPlexExplorer, these networks define principles of proteome organization and enable unknown protein characterization.

## INTRODUCTION

While a cell’s genetic inheritance is fixed, its proteome adapts to external and internal cues, fostering tremendous diversity of cellular form and function that drives multicellular life. Myriad physical interactions assemble proteins into modules that impart spatial and functional organization, thus defining the interactome, an interaction network whose topology encodes each protein’s cellular environment and whose structure may vary with cell state. Defining the full repertoire of protein interactions and the conditions in which they occur will thus be essential to understand proteome diversity.

Despite its importance, a complete map of the human interactome remains elusive due to several challenges: 1) the innumerable proteins, isoforms, and post-translational states within the proteome; 2) the biochemical properties of individual proteins; 3) variable protein expression; 4) the prevalence of transient interactions; and 5) interaction context dependence. Existing interaction profiling methods have only partially addressed these challenges. Binary methods including yeast-two-hybrid assays excel at screening large protein libraries, though interacting protein pairs must be detected in isolation within a foreign cellular environment (Rolland et al., 2014). Alternatively, co-fractionation detects protein interactions in native complexes, subject to limits of sensitivity, dynamic range, and resolution of fractionation (Havugimana et al., 2012; Wan et al., 2015). In contrast, affinity-purification mass spectrometry (AP-MS) enables enrichment and detection of even low-abundance proteins, though extensive sample preparation has limited scalability while precluding recovery of transient interactions (Gingras et al., 2007). Approaches that combine datasets (Drew et al., 2017) or mine literature (Oughtred et al., 2018) can compensate for limitations of individual approaches, though experimental context may be lost (Stacey et al., 2018). Thus, our view of the interactome remains static and fragmentary. While our understanding of interactome dynamics is especially limited, context-dependent interactions enable cells to adapt to variable environments and create the cellular diversity that drives tissue-specific function and disease susceptibility. Such conditional interactions combine with protein expression, localization, and post-translational modification states – the *proteotype* – to biochemically link genotype to phenotype.

Though no single methodology can overcome all limitations, AP-MS excels for profiling interactomes – including yeast (Gavin et al., 2002; Ho et al., 2002; Krogan et al., 2006), *Drosophila* (Guruharsha et al., 2011), and human (Hein et al., 2015) – due to its sensitivity and its ability to detect interactions within complexes in appropriate cellular contexts. Thus, we have established a robust platform for AP-MS profiling of the human interactome, generating BioPlex 1.0 (Huttlin et al., 2015) and BioPlex 2.0 (Huttlin et al., 2017). Here we present BioPlex 3.0, the most complete model of the human interactome to date. This network is a powerful tool for biological discovery whose structure encodes protein function, partitions into communities, and reveals fundamental principles of interactome organization.

Despite its unmatched scale, BioPlex 3.0 depicts just one cellular context. To begin to explore how interactomes vary with cellular state, we have created a second network in HCT116 cells, leading to the first experimentally derived, proteome-scale, cell-specific models of the human interactome. Together, they reveal shared and cell-specific modules with characteristic biological and network properties. In combination, they enable biological discovery, revealing physical interactions and suggesting functions for thousands of human proteins. Both networks are viewable interactively online through BioPlex Explorer.

## RESULTS

### Profiling the Human Interactome at Proteome Scale

Our robust pipeline features lentiviral expression of genes from the human ORFeome v. 8.1 (Yang et al., 2011) in 293T cells (**Figure 1A**). Upon validation each clone is expressed with a C-terminal FLAG-HA tag for enrichment of baits and interactors. HA-purified protein complexes undergo LC-MS after HA peptide elution. Detected proteins are then filtered with *CompPASS-Plus* (Huttlin et al., 2017) to distinguish high-confidence interacting proteins from non-specific background and false positive identifications. Combining thousands of IP’s has produced BioPlex 1.0 (Huttlin et al., 2015) and BioPlex 2.0 (Huttlin et al., 2017) which model the human interactome with increasing coverage (**Figure 1A**). The latest, BioPlex 3.0, incorporates 10,128 AP-MS experiments and results from attempted AP-MS of all validated ORFeome v. 8.1 clones (**Table S1**).

**Figure 1:**
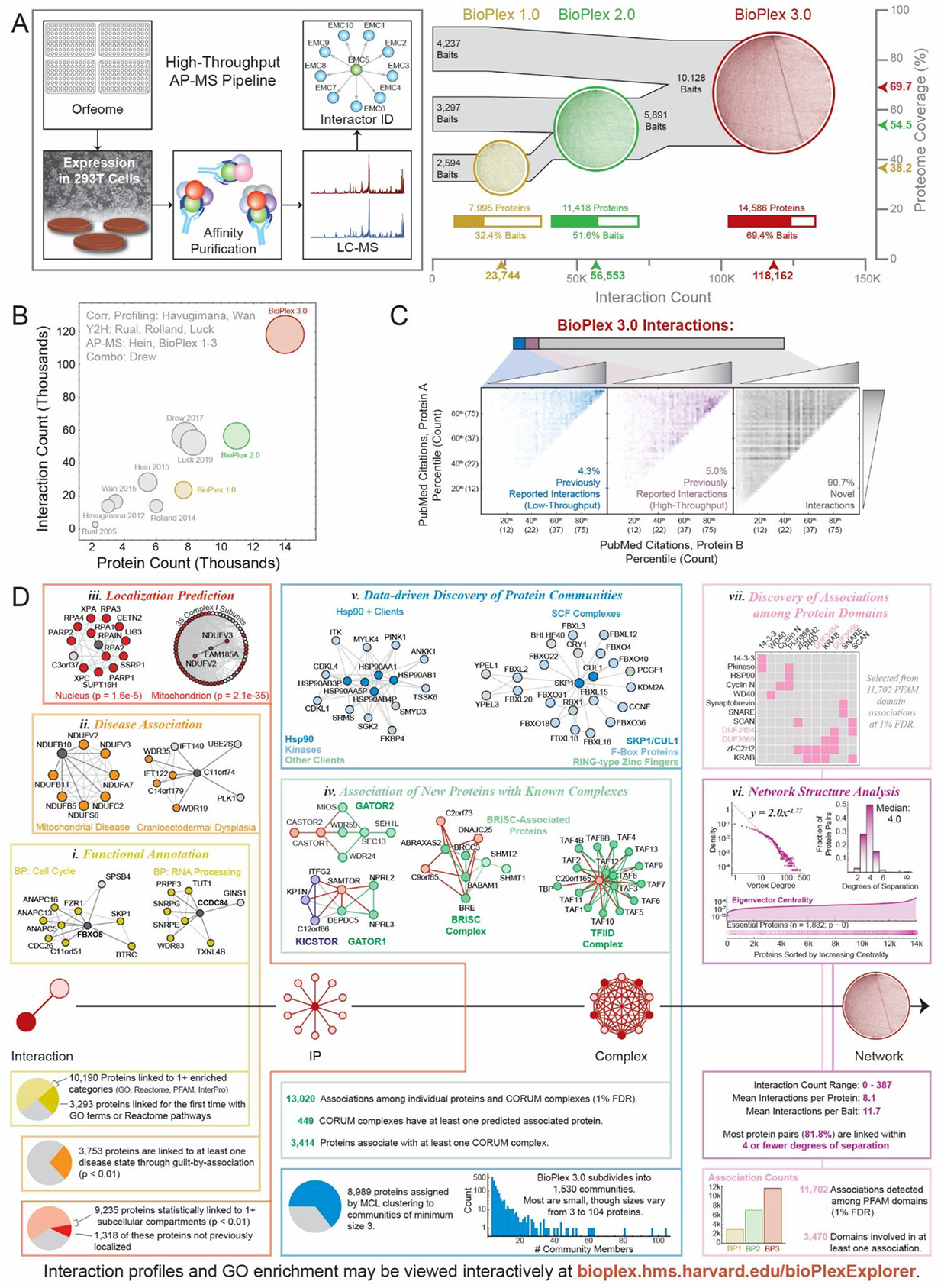
A Proteome-scale Map of the Human Interactome (A) High-throughput interaction profiling via affinity purification-mass spectrometry. See text for details. (B) BioPlex 3.0 expands interactome coverage beyond previous attempts. Y2H: yeast-two-hybrid. See text for citations. (C) Comparison with BioGRID reveals that most BioPlex 3.0 interactions have not been previously reported. Incorporating PubMed citation counts for individual proteins suggests that much of the increased coverage comes from interactions among poorly studied proteins. (D) BioPlex 3.0 enables biological discovery at scales ranging from individual interactions to network-level analyses. See text for details.

BioPlex 3.0 attains proteome scale, encompassing 70% of human proteins. Coverage of some protein classes is higher, including cell fitness genes (90%) (Blomen et al., 2015; Wang et al., 2015), mitochondrial proteins (88%) (Calvo et al., 2016), protein kinases (85%) (kinase.com/web/current/human/), and phosphatases (83%) (Chen et al., 2017). Moreover, BioPlex 3.0 contains most medically significant proteins, including 88% of cancer genes (Vogelstein et al., 2013), 65% of disease genes (Piñero et al., 2017) and 70% of drug targets (www.drugbank.ca). BioPlex 3.0 coverage exceeds previous efforts utilizing yeast-two-hybrid (Luck et al., 2019; Rolland et al., 2014; Rual et al., 2005), correlation-profiling (Havugimana et al., 2012; Wan et al., 2015), AP-MS (Hein et al., 2015; Huttlin et al., 2015, 2017) and combinations thereof (Drew et al., 2017) (**Figure 1B**).

One consequence of its scale is that most BioPlex 3.0 interactions have not been independently reported. Intersecting BioPlex interactions with BioGRID (Oughtred et al., 2018) reveals prior high- or low-throughput detection of just 9% (**Figure 1C**). Binning interactions according to PubMed citations for each protein reveals that those confirmed by BioGRID involve well-studied proteins. In contrast, BioPlex-specific interactions couple well-studied and unknown proteins alike, reflecting the sensitive, unbiased nature of our AP-MS approach.

### Internal and External Validation of BioPlex 3.0

Since most interactions involve minimally characterized proteins with little independent interaction data, validation has required several complementary strategies. To assess coverage of known complexes, BioPlex 3.0 was compared against CORUM (Giurgiu et al., 2018). When a complex is observed in BioPlex, we expect its corresponding subnetwork to be highly interconnected relative to global network density (**Figure S1A**). Indeed, nearly 75% of complexes are so enriched, exceeding prior interaction datasets (**Figure S1B**). This includes prior BioPlex versions; as the network has grown, complex coverage has increased individually (**Figure S1C**) and collectively (**Figure S1B**).

Because CORUM complexes match just a fraction of BioPlex, we also used several additional approaches to validate a larger share of interactions. Since protein complex members at least partially co-elute via size exclusion chromatography, we correlated elution profiles from 293T proteome co-fractionation (Heusel et al., 2019) for all interacting proteins. Elution of BioPlex interacting proteins correlated strongly (**Figure S1D**), suggesting that co-elution confirms many interactions. Similarly, interacting proteins must co-localize. Comparison of subcellular Bar Codes (Orre et al., 2019) revealed that interacting proteins co-purify upon subcellular fractionation (**Figure S1E**). Finally, we measured the assortativity of each GO SLIM category (Ashburner et al., 2000) in BioPlex relative to 1,000 randomized networks (**Figure S1F**). Proteins within nearly all functional categories interacted preferentially with each other.

In addition, experimental confirmation of specific interactions can be obtained from reciprocal AP-MS of interacting protein pairs or by repeatedly purifying a protein complex while targeting different members as baits. As the fraction of baits in BioPlex has increased to 70%, much of the network has become eligible for intra-network confirmation. For reciprocal detection, both interacting proteins must be baits and must appear as preys in 293T cells. Over 40,000 interactions are eligible for reciprocal detection and 29% are detected reciprocally (**Figure S1G**). While technical factors can prevent reciprocal detection of some interactions, the dramatic reciprocal enrichment favors network veracity. These reciprocal interactions include associations among heterodimers (e.g. YWHAG – YWHAB; ERO1L – ERO1LB) and complex members, both direct (e.g. ARPC3 – ACTR3) and indirect (e.g. ARPC3 – ARPC1A; ARPC3 – ARPC5) (Goley and Welch, 2006), as well as pairings involving uncharacterized proteins (e.g. C11orf49 and several polyglutamylase subunits). Analogously, complex co-purification can be assessed via 3-cliques: mutually interacting protein triads (**Figure S1H**). As with reciprocal interactions, 3-cliques are strongly enriched, reflecting modular network architecture and frequent complex co-purification. Half of all edges map to at least one confirmatory 3-clique; well-known examples include SCF complexes (SKP1 – CUL1 – FBXO4), enolase subunits (ENO1/2/3), and katanin subunits (KATNA1/B1/BL1). Including both 3-cliques and reciprocal interactions, most BioPlex edges are confirmed intra-network by at least one complementary IP (**Figure S1I**).

As further validation, we repeated AP-MS of 999 baits in 293T cells (**Figure S1J, Table S2E-F**) and achieved a median 60% replication rate. This compares favorably with previous reports (Varjosalo et al., 2013), especially since these experiments were performed without individual bait optimization. Replication profiles of several baits are displayed; high overlap was seen for many, including CASTOR1 (100%), C12orf34 (100%), and CDC20 (89%). When replication was lower, other BioPlex interactions often supported non-replicated edges. Examples include C12orf10 (25%), whose three non-replicated interactions are supported by pulldown of NOTCH2HL; similarly, though REG1B pulldown of REG1A was not replicated, the interaction has been confirmed reciprocally.

### BioPlex 3.0 Enables Biological Discovery

As we have described previously (Huttlin et al., 2015, 2017), diverse insights arise from viewing BioPlex at multiple scales (**Figure 1D**). Locally, a protein’s neighbors can reveal localization, function, and disease associations via guilt-by-association. When GO, Reactome (Fabregat et al., 2017), PFAM (El-Gebali et al., 2018), and InterPro (Mitchell et al., 2018) gene sets were mapped to BioPlex 3.0, 10,190 proteins associated with at least one enriched gene set, including FBXO5 (also called EMI1), an anaphase-promoting complex regulator whose neighbors are enriched for the GO Biological Process term “Cell Cycle” (**Figure 1D*i***, **Table S1C-H**). Notably, 3,293 of these proteins were not previously linked to GO or Reactome terms, including CCDC84, whose neighbors suggest a role in RNA processing. Similarly, layering DisGeNET (Piñero et al., 2017) onto BioPlex links 3,753 proteins to at least one disease (**Figure 1D*ii***, **Table S1I**). For example, C11orf74 was recently confirmed to interact with the IFT-A complex (Takahara et al., 2018), whose members are associated with Cranioectodermal Dysplasia. Finally, subcellular localization may be inferred for 9,235 proteins, including FAM185A and RPAIN, from Uniprot localizations of their primary and secondary neighbors; 14% have no prior known localization (**Figure 1D*iii***, **Table S1J**).

BioPlex can also associate new proteins with known complexes (**Figure 1D*iv***). Highlights include CASTOR1/2 (née GATSL3/2), which interact with the GATOR2 complex and monitor cytosolic arginine (Chantranupong et al., 2016), as well as SAMTOR (née C7orf60) which binds to GATOR1 and KICSTOR complexes and senses S-adenosyl-methionine (Gu et al., 2017). Systematic analysis of CORUM complexes reveals 13,020 associations linking 3,414 proteins with 449 complexes; examples include DNAJC25, C2orf73, and C9orf85 and the BRISC complex as well as C20orf65 and the TFIID complex. We have also discovered protein communities using network structure without prior biological knowledge (**Figure 1D*v***, **Table S1K-L**). MCL clustering (Enright et al., 2002) subdivides BioPlex 3.0 into 1,590 communities ranging from 3 to 104 proteins. Though defined from network connectivity alone, most correlate with function, as shown for HSP90 subunits with kinases and other clients as well as CUL1, SKP1, and assorted F-box proteins.

BioPlex also reveals principles of interactome organization (**Figure 1D*vi***). The average protein participates in 8.1 interactions, though this varies dramatically. The network degree distribution approximates a power law, though deviations at low counts suggest that proteins with few interacting partners are disfavored. The interactome is dense, with 82% of proteins linked by 4 or fewer degrees of separation. Moreover, ‘essential’ proteins occupy the most central network positions. Additionally, enrichment analysis reveals PFAM domain pairs that associate directly or indirectly (**Figure 1D*vii***, **Table S1M**). Examples include associations among HSP90 and protein kinase domains; SNARES and synaptobrevins; as well as self-associations among protein kinase domains, KRAB domains, and 14-3-3 domains. Some associations involve domains of unknown function: previously we reported association of DUF3669 and KRAB domains (Huttlin et al., 2017); DUF3669 has since been shown to trigger interactions among KRAB-domain-containing zinc finger proteins (Helleboid et al., 2019).

### The Protein Interaction Network of a Second Human Cell Line

While BioPlex 3.0 enables biological discovery, it samples one cell type under standardized growth conditions and thus only partially models the dynamic interactome. As a first step toward understanding interactome diversity, we have created a second interaction network in HCT116 cells.

Though both 293T and HCT116 cells grow robustly and readily express exogenous proteins, suiting them for large-scale AP-MS, they differ in sex, tissue of origin, karyotype, and driver modifications (**Figure S2A**). To explore these differences, we performed RTS-MS3-TMT quantitative proteomics (Erickson et al., 2019) and found that 54% of proteins are differentially expressed (**Figure S2A, Table S2A**), reflecting the distinct origin of each cell line. While 239T-specific proteins were enriched for embryonic/nervous system development, evoking their embryonic kidney/adrenal origin (Stepanenko and Dmitrenko, 2015), proteins specific to colorectal-cancer-derived HCT116 cells were enriched for cell adhesion and cadherin binding. In particular, marker proteins reflect the male status of HCT116 cells (EIF1AY), the potential for 293T cells to ciliate (CEP290) (Takahashi et al., 2018) and the contrasting epithelial (GRHL2, CDH1, LAMC2) and mesenchymal (TBX2, CDH2, VIM) origins of HCT116 and 293T cells (**Figure S2B**). In sum, phenotypic and proteotypic differences ensure that these cell-specific networks will provide contrasting views of the human interactome.

Because only 5,522 of 10,128 successful BioPlex 3.0 AP-MS experiments have been repeated in HCT116 cells, this network is smaller than BioPlex 3.0 (**Figure 2A, Table S2B-C**). Nevertheless, aside from BioPlex 3.0, our HCT116 network exceeds all other experimentally derived interaction networks in size (**Figure 2B**). Both networks contain similar proteins, with 67% shared (**Figure 2C**). While 4,463 proteins are specific to the 293T network, 72% are baits that have not yet been targeted in HCT116 cells. In contrast, only 408 proteins are unique to HCT116 cells, and most are preys. Inset histograms indicate that differential expression explains only a subset of cell-line-specific proteins (**Figure 2C**). Subnetworks surrounding HCT116-specific proteins are shown (**Figure 2E-H**). For RIOX1, ∼5-fold higher expression in HCT116 cells may explain cell-specific detection; in contrast, comparable expression (∼1.2-fold) suggests that cell-specific detection of MRPL54 is driven by other factors.

**Figure 2:**
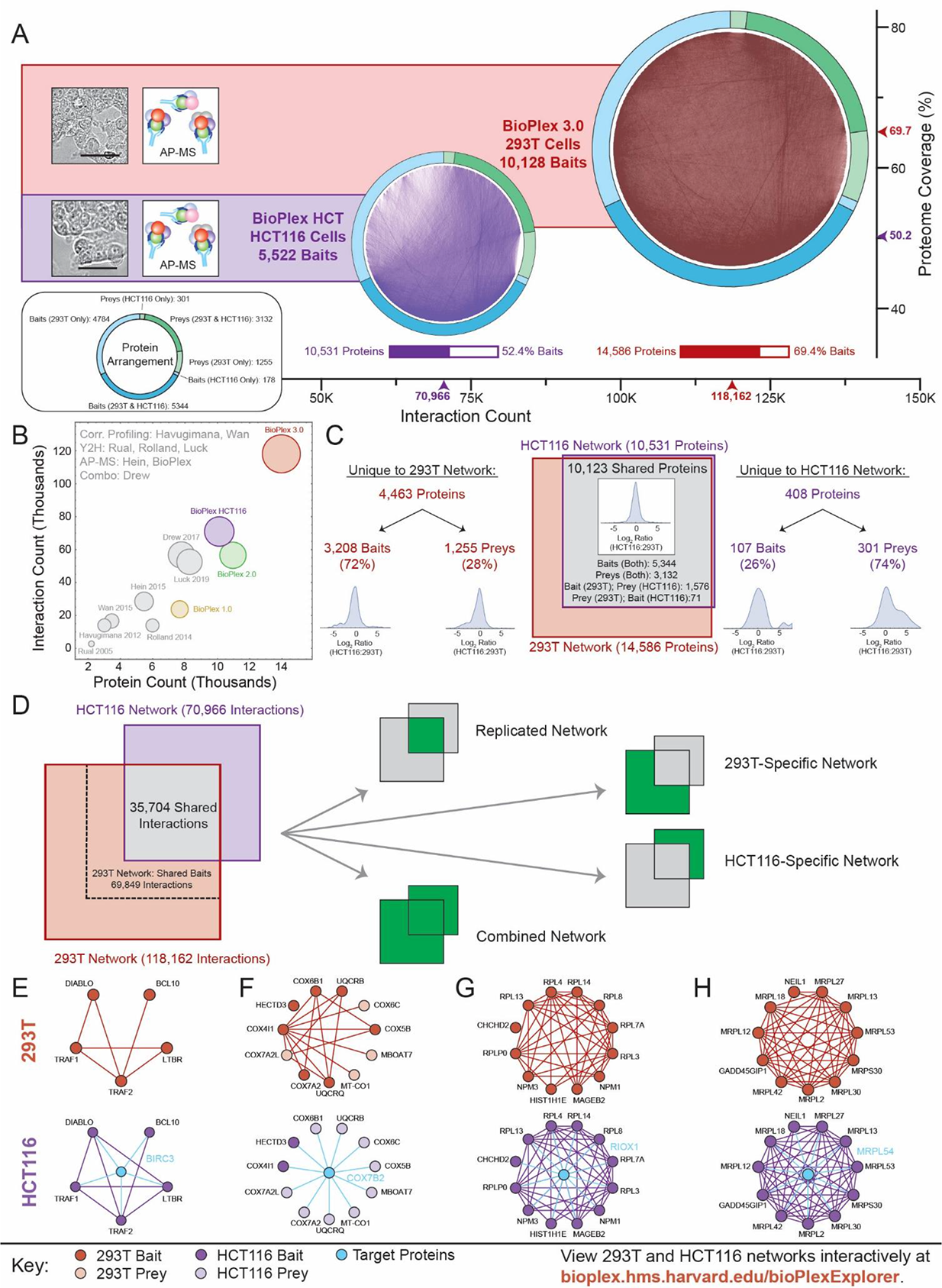
A Second Proteome-scale, Cell-type-specific Human Interaction Network. (A) We repeated AP-MS analysis of ORFeome baits in HCT116 cells to produce a second interaction network (BioPlexHCT 1.0) that complements our 293T network (BioPlex 3.0). (B) Aside from BioPlex 3.0, BioPlexHCT 1.0 already exceeds all prior experimentally derived human interaction networks in depth and breadth. (C) Overlap among proteins in 293T and HCT116 interaction networks. Inset histograms depict Log_2_ protein expression ratios between cell lines (see **Figure S2**). (D) Overlap among interactions in 293T and HCT116 interaction networks. A dashed box depicts the subset of 293T interactions through AP-MS analysis of baits that have also been targeted in HCT116 cells. (E) – (H) Sub-networks surrounding HCT116-specific target proteins BIRC3, COX7B2, RIOX1, and MRPL54.

Although 293T and HCT116 networks share most proteins, overlap among interactions is modest, with 35,704 interactions shared (**Figure 2D, Table S2D**). After removing IP’s not repeated in HCT116 cells, this represents ∼50% overlap among remaining interactions. Partitioning interactions according to their detection in each cell line defines complementary subnetworks: their union defines a combined network with maximal coverage; in contrast, their intersection defines a “Replicated Network” whose edges have been observed multiple times in distinct cell lines and are thus prime targets for biological study. Finally, cell-specific edges may reflect proteotypic differences.

### Interaction Overlap Reflects Protein Abundance, Essentiality, and Network Context

To understand how cell context influenced interaction replication, we repeated AP-MS of selected clones within the same cell line or across cell lines. 999 AP-MS experiments were repeated in 293T cells (**Figure S1C, Table S2E-F**) and 72 in HCT116 cells (**Table S2G-H**), with 5,112 AP-MS experiments replicated across cell lines (**Figure S2C, Table S2I-J**). Median clone-wise reproducibility rates nearly doubled within the same cell line (**Figure S2D**), suggesting that technical and biological factors linked to cellular context account for much inter-network variability.

Bait and prey expression contribute to AP-MS technical variability. Since most proteins are differentially expressed between cell lines (**Figure S2A**), prey abundance will vary. Bait abundance also varies, with HCT116 cells achieving 2-fold lower bait expression (**Figure S2E**). Decreases in either bait or prey abundance reduce interaction replication in HCT116 cells (**Figure S2F-G**). Subnetworks surrounding 293T-specific proteins CDH2 and MYEF2 as well as HCT116-specific proteins FAM111B and RAC2 show how protein differential expression can account for some cell-specific interactions (**Figure S2H-K**).

One interpretation of this is biophysical: decreased bait or prey levels can reduce the likelihood of interaction while making detection more difficult. Variations in exogenous bait expression thus directly affect reproducibility. In contrast, although variable prey abundance exerts an analogous effect, a second possibility is that both prey abundance and interactions vary due to differential cell biology. In fact, several lines of evidence suggest that shared and cell-line-specific interactions differ in network context and biology.

To assess properties of shared and cell-specific interactions, 293T and HCT116 networks were combined and edges scored for betweenness centrality and local clustering coefficient. The fraction of edges shared between cell lines varied inversely with edge betweenness centrality (**Figure S3A**) and jointly with local clustering coefficient (**Figure S3B**), suggesting that shared interactions reside in dense subnetworks while cell-specific interactions bridge otherwise disparate proteins and complexes, effectively “rewiring” connections among core modules to suit each cell’s needs.

We next assessed how cell fitness and protein expression variability influence interaction overlap. Superimposing cell fitness data (Blomen et al., 2015; Dempster et al., 2019; Wang et al., 2015) onto the combined BioPlex network revealed a strong positive correlation between “essential” proteins and interaction overlap (**Figure S3C**). Similarly, ranking by expression variability across cancer cell lines (Nusinow et al. *in press*) (**Figure S3D**) or human tissues (Wang et al., 2019) (**Figure S3E/F**) indicates that proteins with cell-specific interaction profiles more generally tend to be variably expressed across diverse biological contexts. Finally, cell-specific interactions disproportionately involve poorly-characterized proteins (**Figure S3G**), highlighting the importance of global, untargeted strategies to define cell-specific interactomes.

### Interaction Overlap Reflects Protein Evolution

We next mapped protein evolutionary ages (Liebeskind et al., 2016) onto BioPlex networks to evaluate the evolutionary context of cell line specificity (**Figure S4A**). Assigning each interaction the age of its youngest constituent protein split the interactome into subnetworks traceable to eight evolutionary stages (**Figure S4B**). While some proteins and interactions dated to the dawn of cellular life, most arose during eukaryotic and eumetazoan ages. A striking correlation was observed between cell line specificity and evolutionary age with the “oldest” interactions overlapping ∼6-fold more often than their “youngest” counterparts (**Figure S4C/D**).

This relationship between shared interactions and evolutionary age also appears in individual protein interaction profiles. Among DDX31 interactors, cell-specific replication drops from 100% to 35% as younger proteins are added (**Figure S4E**). Similar trends are observed for HDAC1 (**Figure S4F**), LIG3 (**Figure S4H**) and the uncharacterized protein C15orf41 (**Figure S4G**). While cell-specific interactions of these proteins favor 293T cells, the GPR180 subnetwork is strongly HCT116-specific (**Figure S4I**) and the CCDC12 subnetwork (**Figure S4J**) is relatively balanced.

### Functionally Defined Subnetworks are Shared Variably across Cell Lines

To relate cell line specificity with interactome functional organization, we compared HCT116- and 293T-specific subnetworks corresponding to Reactome pathways. Though intra-pathway overlap ranged from 0 to 100%, most exceeded 75% (**Figure 3A**). Pathways with highest overlap match fundamental cellular processes, including glycolysis (**Figure 3D**), for which virtually every interaction is observed in both cell lines after accounting for baits targeted in each. In contrast, cell-specific subnetworks denote pathways relevant to each cell’s unique biology. For example, interactions among EPH-Ephrin signaling proteins are predominantly detected in 293T cells (**Figure 3B**). Ephrins and ephrin receptors are cell surface proteins whose binding triggers complementary signaling cascades in neighboring cells, contributing variously to cell migration, repulsion, and adhesion in various cellular contexts, especially during development (Kania and Klein, 2016). Still more striking are cell-specific interactions within the pathway “Transcription of E2F Targets under Negative Control by RBL1 and RBL2 in complex with HDAC1”: in HCT116 cells these proteins form the DREAM (DP, RB-like, E2F, and MuvB) complex, whose structure disintegrates in 293T cells (**Figure 3C**). The DREAM complex silences cell cycle genes during G0. Unlike HCT116, 293T cells express SV40 large T antigen that binds and sequesters RBL1/2, blocking DREAM and RB-E2F complex formation, favoring cell cycle gene expression and reducing growth inhibition to promote cell line immortality (Sadasivam and DeCaprio, 2013). Additional Reactome pathways may be viewed through BioPlexExplorer.

**Figure 3:**
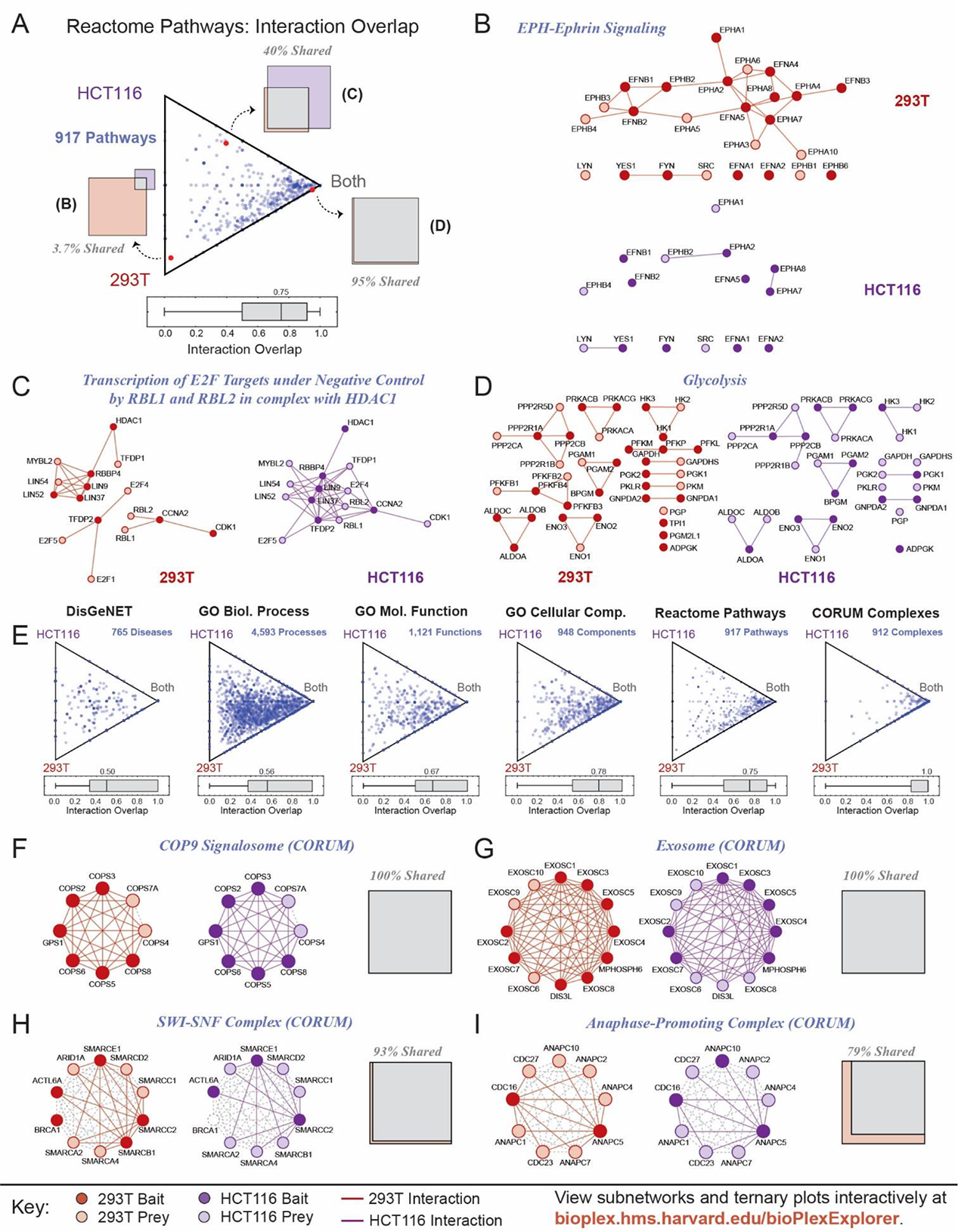
Functionally-defined Subnetworks are Shared Variably across Cell Lines (A) Ternary diagram depicting the proportions of edges shared or unique to either 293T or HCT116 cells for subnetworks defined by each of 917 Reactome pathways. Proteins in each pathway were mapped to the combined 293T/HCT116 network to define a subnetwork whose edges were shared or unique to either cell line. Individual pathways are displayed as Venn diagrams as shown for 3 examples; each may also be represented as a point within the ternary diagram whose location reflects the relative proportions of shared and cell-specific edges. Points near the corners indicate that most edges are either shared (“Both”) or cell-specific; points near the center of the triangle indicate edges evenly distributed across shared and cell-specific categories. The ternary diagram thus summarizes Venn diagrams for 917 pathways. A box-whisker plot depicts the edge overlap across pathways. (B) – (D) Subnetworks corresponding to three pathways highlighted in panel A. (E) Ternary diagrams depicting edge sharing across cell lines for subnetworks defined by protein functional classifications. Separate plots are provided for CORUM complexes, Reactome pathways, GO ontologies, and DisGeNET disease associations, ordered by edge sharing among constituent protein classes. (F) – (I) Subnetworks corresponding to four CORUM complexes. Venn diagrams accompany each complex.

We next extracted 293T and HCT116 subnetworks matching CORUM complexes, GO categories, and DisGeNET disease associations and quantified overlap within each (**Table S3**). Cell-line specificity varied with category (**Figure 3E**); those most closely associated with physical entities (e.g. CORUM, GO cellular components) overlapped more than abstract functional classes (e.g. GO biological processes, disease associations). Since most perform essential functions, CORUM complexes were most consistent, with over 50% interacting identically in 293T and HCT116 cells. Examples include the COP9 Signalosome, Exosome, SWI-SNF Complex, and Anaphase-promoting Complex (**Figure 3F-I**). In contrast, greater cell specificity observed for GO categories and DisGeNET disease associations reflects specialization of individual cells and the tissue-specific nature of disease. For example, interaction networks associated with pre- and post-synaptic membranes or delayed rectifier K+ channel activity (**Figure S5A-C**) form exclusively in 293T cells. 293T cells can be induced to form pre- and post-synaptic structures (Biederer and Scheiffele, 2007), suggesting that these interactions reflect 293T cells’ neural crest derivation (Stepanenko and Dmitrenko, 2015). Similarly, HCT116-specific interactions among pyrimidine processing proteins likely reflect unique metabolism (**Figure S5D**). Finally, reorganization of extracellular matrix protein interactions (**Figure S5E**) may reflect distinct surface protein organization in epithelial and mesenchymal cells.

### Data-driven Discovery of Shared and Cell-line-specific Network Communities

Mapping CORUM complexes, Reactome Pathways, GO Categories, and DisGeNET associations onto HCT116 and 293T networks has leveraged prior knowledge to link cell-specific interactions with known cellular structures and protein functions. As an alternative, data-driven approach, we used MCL clustering to discover protein communities within the combined 293T/HCT116 network (**Table S4A-B**). We then sought statistical associations among community pairs whose members were especially likely to interact (**Table S4C**) and quantified interaction overlap within and between communities (**Figure 4A, Table S4D-E**). Together, these communities and associations define a network displayed in **Figure 4B** and on BioPlexExplorer. Because network architecture defines these communities and associations independently of existing biological knowledge, they model the interactome more completely, incorporating characterized and uncharacterized proteins and complexes without knowledge bias.

**Figure 4:**
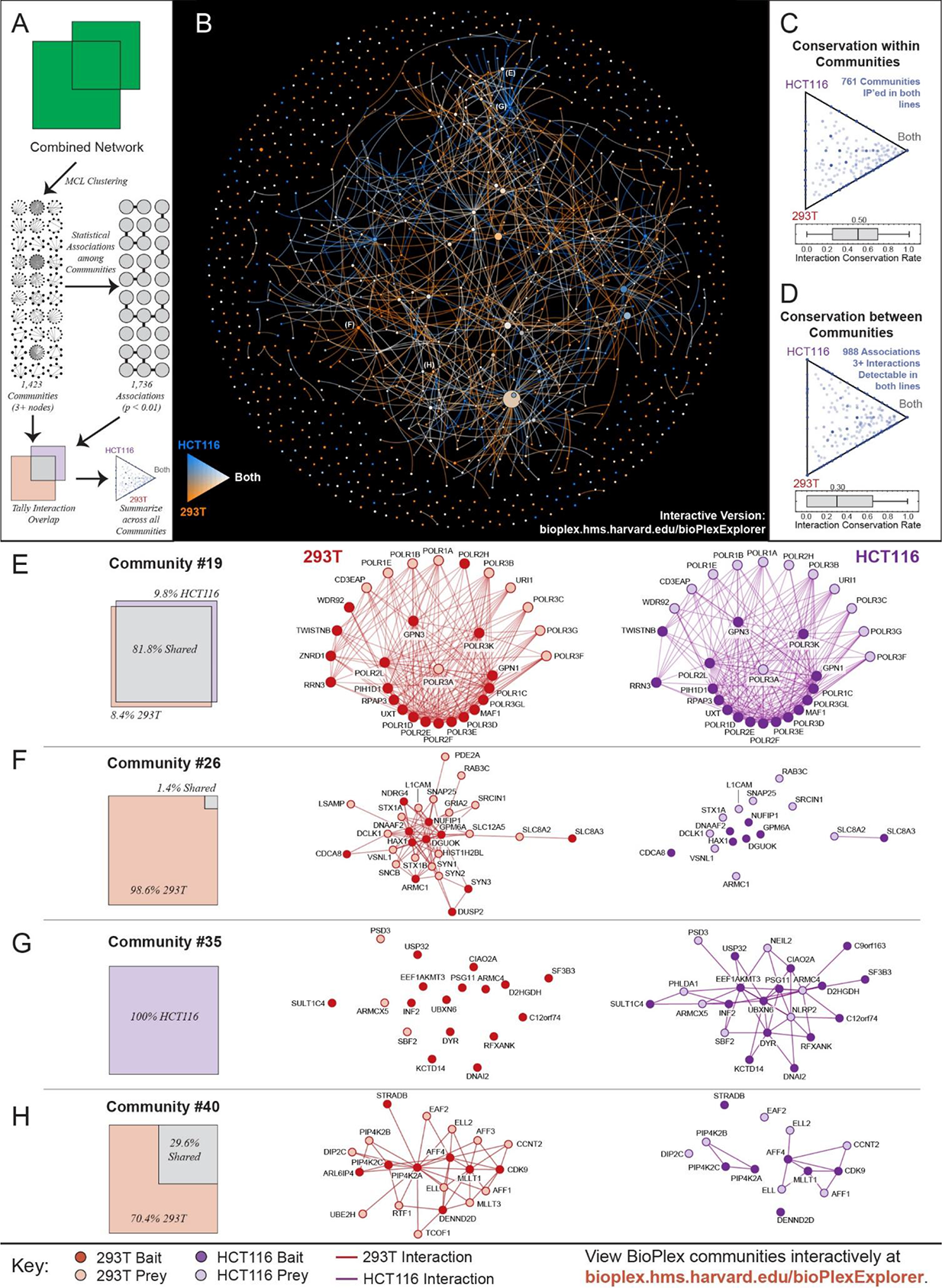
Data-driven Discovery of Shared and Cell-line-specific Network Communities (A) 293T and HCT116 networks were combined and partitioned via MCL clustering to identify 1,423 network communities with 3+ members. Interactions among communities were then tallied to identify 1,736 statistically enriched associations. Interaction overlap across cell types was then tallied within each community and along edges that connect associated community pairs. (B) Network of communities detected in the combined 293T/HCT116 interaction network. Every network community with at least 3 members is represented as a node whose size is proportional to the number of proteins it contains. Edges connect communities that were statistically associated. Nodes and colors reflect the level of overlap among cell lines. (C) Ternary diagram depicting the overlap observed within each community. Only communities for which at least one member has been a bait in both cell lines are included. (D) Ternary diagram depicting the overlap observed for edges that connect communities. Edges were only included if they were supported by at least 3 edges detectable in both cell lines given the baits targeted in each. (E) – (H) Selected Network Communities.

Analysis of interaction overlap within communities (**Figure 4C**) and community pairs (**Figure 4D**) reveals both shared and cell-specific interactions. Overlap within communities significantly exceeds overlap between communities, implying multi-tiered interactome organization where shared complexes interconnect in cell-specific ways. Compared to knowledge-driven protein functional classes (**Figure 3E**), overlap among edges within and between data-driven communities is remarkably low. The median 50% edge overlap within these communities trails all literature-derived classifications – especially CORUM and GO Cellular Component categories that most directly encapsulate known protein complexes. This suggests that our current knowledge is biased and incomplete, focusing disproportionately on universal complexes that represent just a fraction of the interactome. Indeed, while some data-driven communities match known protein complexes, most of whose interactions are detected in both cell lines (**Figure 4E**), other communities are exclusively detected in either cell line (**Figure 4F-G**). While biological roles of these latter two complexes are uncertain, their cell-specific co-purification with many community members implies context-dependent assembly. Finally, partial interaction overlap within a community can reveal cell-specific processes. In 293T cells we detect nuclear proteins bound to multiple PIP4K2 subunits; 70% of these interactions are undetectable in HCT116 cells, separating PIP4K2 subunits from nuclear proteins (**Figure 4H**). Since PIP4K2 subunits localize to multiple cellular compartments, these 293T-specific interactions could suggest coordinated nuclear translocation.

### Though Interactions Differ across Cell Lines, they are Functionally Consistent

Given the considerable variation observed, a natural question is whether cell-specific interactions of individual proteins share common biological functions. We thus analyzed the enrichment of functional terms among each bait’s neighbors in the combined BioPlex network. Proteins matching each enriched term were categorized according to detection in one or both cell lines and each protein – function association plotted to reflect its overlap between cell lines (**Figure S6A**). For ECHDC2, only 4/10 interacting proteins were observed in both cell lines; yet nearly all are mitochondrial and several link to mitochondrial organization and branched-chain amino acid metabolism. When this analysis is extended to all baits targeted in both cell lines, 90% of protein – function associations map to both (**Figure S6B, Table S5C**).

Though a protein’s neighbors may differ across cell lines, they often share biological function. This is demonstrated by TRIM28, a protein known to recruit gene-silencing complexes to specific genomic regions upon binding to KRAB-domain-containing zinc-finger proteins (**Figure S6C**). As expected, 92% of its 157 neighbors contain KRAB domains. Only 80 TRIM28 interactors are shared; 55 are HCT116-specific and 22 were detected in 293T only. Differential protein expression explains only a fraction of these cell-specific interactions. That TRIM28 would bind different zinc-finger-containing proteins in 293T and HCT116 cells is not unexpected, as gene silencing differs in each. Similarly, we find that the phosphatase inhibitor FAM122A binds distinct phosphatases (**Figure S6D**) while Kelch/BTP protein KLHL29 binds cell-specific subsets of BTB proteins (**Figure S6E**) and TWIST2 binds proteins associated with DNA-templated transcription (**Figure S6F**). Finally, many uncharacterized proteins have functionally consistent, yet context-dependent interaction profiles (**Figure S6G-J**).

### Cell-line Specificity among Domain Associations

Beyond reporting specific interactions, the BioPlex networks also encode recurring interactions among protein families, as previously shown for domain-domain associations (**Figure 1Dvii**). To understand how domain-domain associations vary with cell type, we mapped PFAM domains onto the combined BioPlex network and identified pairs of protein domain families whose members are likely to interact (**Table S5A**). We then extracted interactions linking each domain pair and calculated their overlap in both cell lines (**Figure 5A, Table S5B**). The resulting domain associations define a network viewable through BioPlexExplorer. Though interactions vary, with a median 40% overlap, most associations (88%) are detected in both cell lines, suggesting that fundamental patterns of protein interaction transcend cell type.

**Figure 5:**
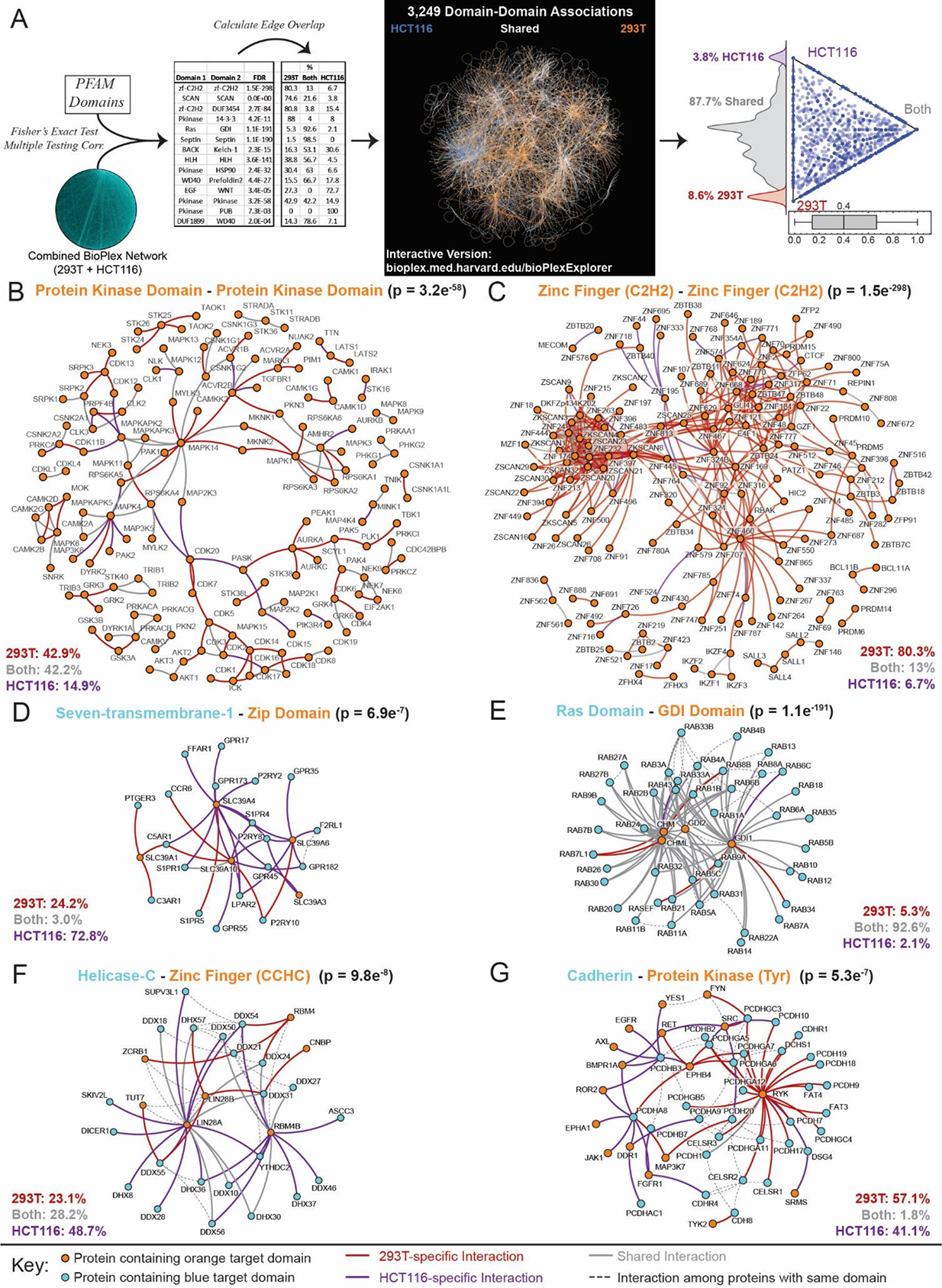
Domain-domain Associations across Cell Lines. (A) PFAM domains were mapped to all proteins in the combined HCT116/293T network. Domain pairs connected by unusually high numbers of edges were then identified. The overlap of edges connecting each statistically associated domain pair was then determined across cell lines. These domain-domain associations are depicted as a network with edge colors that reflect sharing of interactions across cell lines. Sharing across cell lines of the interactions underlying each domain association is also depicted as a ternary plot. The box-whisker plot shows the fraction of interactions shared among cell lines while a histogram highlights the share of domain associations that are either shared or cell-line-specific. (B) – (G) Subnetworks of associated PFAM domain pairs. P-values reflect enrichment of interactions among the indicated domain pair, following multiple testing correction. Only edges eligible for detection in both cell lines are shown.

While most domain associations appear in both cell types, the underlying interactions often reveal stark cell-specific differences. One example is the kinase domain family (**Figure 5B**). Kinases partition into clusters corresponding to CDK’s, MAP kinases, and others. After removing AP-MS experiments not performed in both cell lines, few HCT116-specific edges are seen, though 42% appear only in 293T cells and suggest cell-specific signaling. Another example is the C2H2-zinc-finger family. These proteins often bind DNA via C2H2 zinc fingers and dimerize, frequently through SCAN or BTB domains (**Figure 5C**). More than 80% of these interactions are detected only in 293T cells, though the basis for this selectivity is unclear. In contrast, interactions among Zip domain-containing proteins and 7-transmembrane-domain-containing proteins strongly favor HCT116 cells (**Figure 5D**), suggesting specialized zinc transport, while Ras-GDI interactions are shared almost entirely between cell types (**Figure 5E**). Moreover, CCHC-zinc-finger-Helicase-C domain interactions occur predominantly in HCT116 cells (**Figure 5F**) while distinct interactions among cadherins and tyrosine kinases are observed in both 293T and HCT116 cells (**Figure 5G**), suggesting cell surface proteome reorganization.

### BioPlex Networks Enable Association of Accessory Proteins with Protein Complexes

Since our 293T and HCT116 networks afford complementary views of the interactome, analyzing them in tandem may improve their predictive power. Previously, we and others have shown that individual interaction networks can predict a protein’s function, subcellular localization, and disease associations (Drew et al., 2017; Huttlin et al., 2015, 2017; Luck et al., 2019; Menche et al., 2015). Combining predictions from two independent networks should enhance detection of true positives and minimize false positives.

As a demonstration, we sought to associate new interactors to known complexes within our networks. Since core complexes show high overlap among cell lines, accessory proteins are likely to be consistently detected. In this context, accessory proteins could include low-stoichiometry proteins that engender an alternative function to a defined complex or proteins that link two complexes of distinct function. We thus scored the enrichment of edges linking neighboring proteins to target complexes in each network independently and then combined p-values to derive final predictions (**Figure 6A**). CORUM complexes detected in the BioPlex networks were targeted for leave-one-out cross-validation and for identification of accessory proteins (**Figure 6B, Table S6A-B**). Cross-validation revealed that using both networks out-performed either network individually. Moreover, 96% of targeted complexes matched at least one accessory protein, with over 16,000 associations predicted.

**Figure 6:**
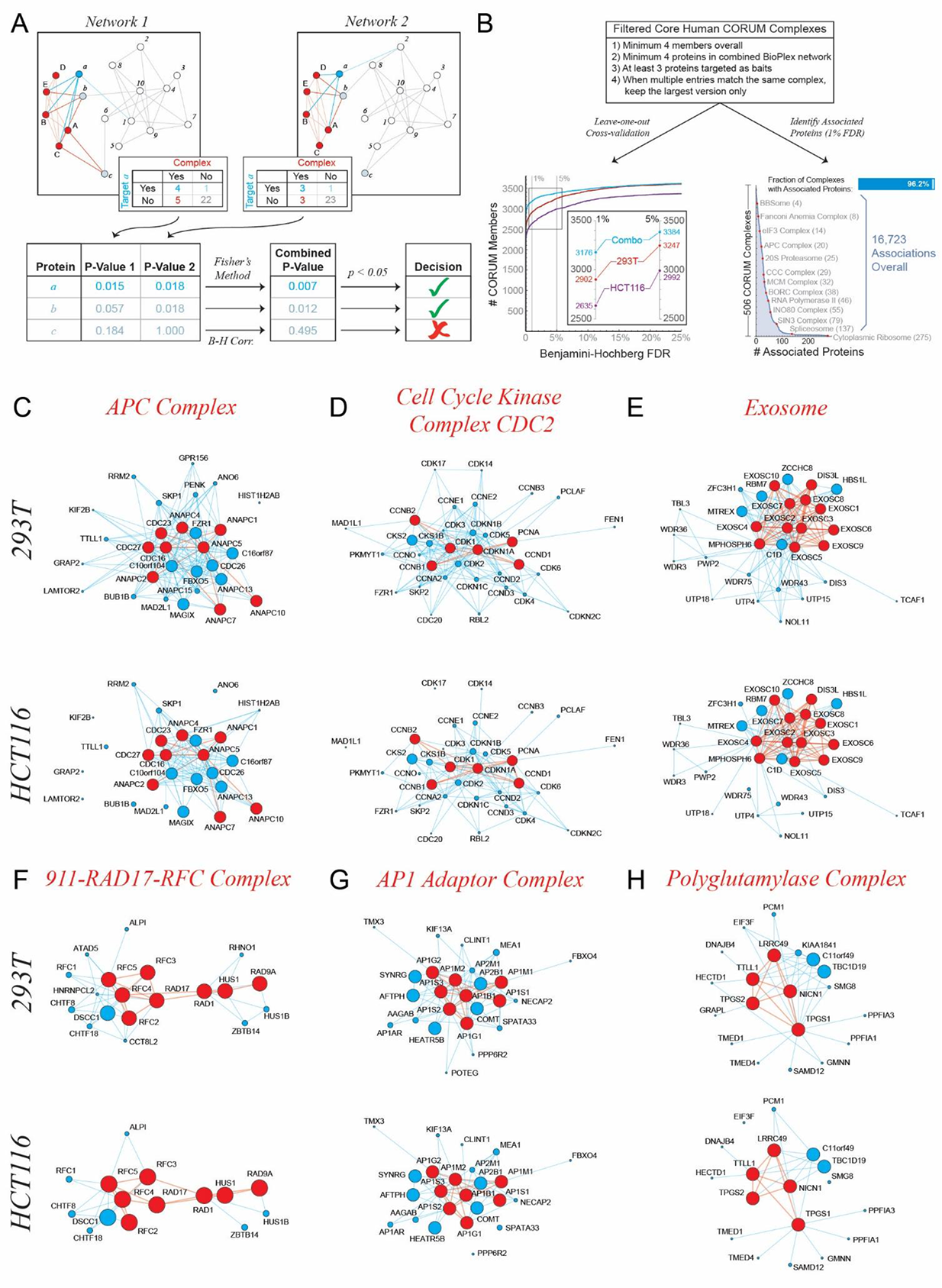
Assignment of Accessory Proteins to Known Complexes. (A) 293T and HCT116 networks associate proteins with known complexes. Within a single network, proteins belonging to a complex are identified (red) along with their neighbors (blue). The enrichment of edges linking each target protein to the complex is scored using Fisher’s Exact Test and adjusted for multiple testing to identify associated proteins. P-values may be combined via Fisher’s Method to integrate results across networks prior to multiple testing correction. (B) CORUM complexes were selected with sufficient coverage in both networks to facilitate leave-one-out cross-validation. 293T and HCT116 networks were used individually and collectively to recover CORUM complex members and compared at multiple false discovery rates (lefthand plot). We further sought additional proteins associated with these complexes, identifying accessory proteins for over 96% of tested complexes (righthand plot). (C) – (H) Each panel presents a single known protein complex along with accessory proteins identified from interactions in 293T and HCT116 networks. CORUM complex members are red, while accessory proteins are colored blue and scaled according to the confidence of the association (larger nodes are more confident).

Predictions for several complexes are displayed in **Figure 6C-H**. While some CORUM complex definitions appeared highly restrictive, excluding known complex members, our predictions were nonetheless robust to incomplete complexes, simultaneously recovering known interactions not annotated in CORUM and identifying novel associated proteins. Among APC neighbors, several additional subunits and associated proteins were recovered (**Figure 6C**), including CDC26, ANAPC13, ANAPC15, FZR1, FBXO5, MAD2L1, and BUB1B in addition to uncharacterized proteins C16orf87, MAGIX, and C10orf104. Similarly, the Cell Cycle Kinase Complex CDC2 associated with many CDK’s and cyclins (**Figure 6D**) and nucleolar proteins surrounded the Exosome (**Figure 6E**), while 911-RAD17-RFC and AP1 complexes linked to known and unknown interactors (**Figure 6F-G**).

For the TTLL1 tubulin polyglutamylase complex, we predicted association of uncharacterized proteins C11orf49, TBC1D19, and KIAA1841 (**Figure 6H, Table S6C**). Because C11orf49 was detected in both cell lines and confirmed by many reciprocal interactions (**Figure S1G**), we selected it for validation, demonstrating binding of endogenous C11orf49 to other polyglutamylase complex members (**Figure S7A,B**). Two distinct C11orf49 isoforms localized to the pericentriolar region, along with established polyglutamylase member LRRC49 (**Figure S7C-E**), in the vicinity of gamma-tubulin (**Figure S7F**).

### Linking Physical and Functional Associations for Biological Discovery

Individually and collectively, both 293T and HCT116 networks afford myriad biological insights. Nevertheless, incorporating complementary datasets can further enable biological discovery. Thus, we combined BioPlex networks with genome-wide CRISPR co-essentiality profiles measured across hundreds of cancer cell lines through Project Achilles (Meyers et al., 2017). Previously, covarying fitness effects have revealed functional associations within protein complexes (Boyle et al., 2018; Pan et al., 2018) and identified novel complex members (Wainberg et al., 2019). We thus correlated fitness profiles for each interacting protein pair in the combined BioPlex network (**Figure 7A**, **Table S7**) and extracted subnetworks corresponding to interactions with either positive (**Figure 7B**) or negative (**Figure 7C**) correlations. While the cell specificity of negatively correlated edges matched the background distribution of edges for which fitness profiles were available, positively correlated edges were enriched for shared edges. This reflects a key structural difference: whereas the positive network includes numerous complexes, the negative correlation network contains none. This is driven by the mathematical impossibility of 3 or more proteins correlating negatively with each other. While entire complexes may correlate positively with each other– and will have higher probability of conservation across cell lines – negative correlations connect specific members of individual complexes or span multiple complexes. Thus, positive and negative correlations reflect fundamentally different functional and structural relationships.

**Figure 7:**
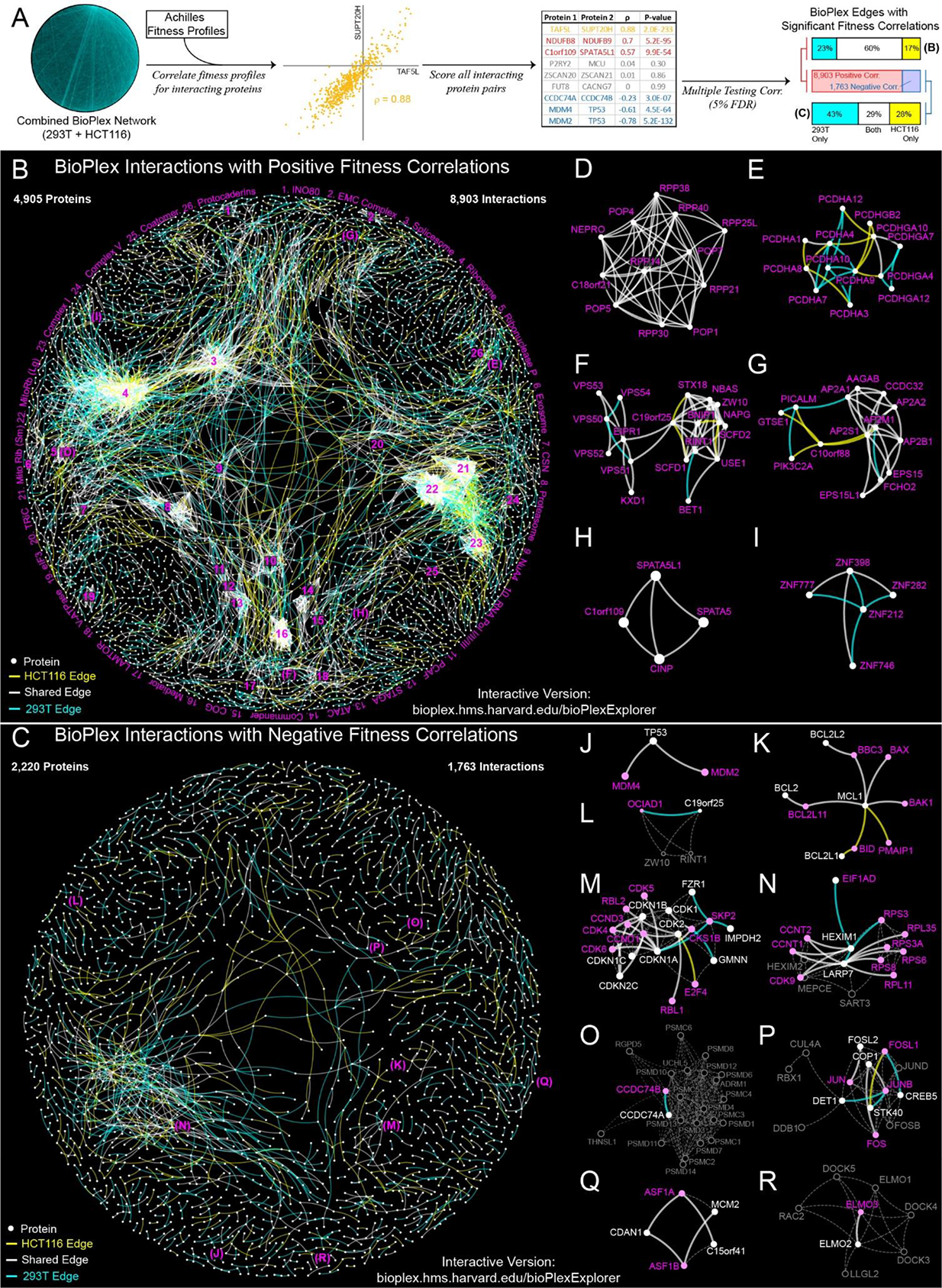
BioPlex and Achilles: Linking Physical and Functional Associations for Biological Discovery. (A) For each pair of interacting proteins in the combined 293T/HCT116 interaction network, cellular fitness profiles from Project Achilles were correlated and assessed for statistical significance. Following multiple testing correction, edges with either positive or negative cellular fitness correlations were extracted and assigned as either shared or cell-specific. Only edges detectable in both cell lines are shown. (B) – (C) BioPlex subnetworks displaying positive **(B)** or negative **(C)** fitness correlations. Edges connect proteins that interact in BioPlex and whose fitness profiles correlate across ∼600 cell lines (5% FDR). Magenta numbers identify selected protein complexes **(B)**. (D) – (I) Subnetworks of panel **B**. Colors match panel **B**. (J) – (R) Subnetworks of panel **C**. Edge colors match panel **C**. Nodes and labels alternate between white and pink to accentuate alternating negative correlations. Other proteins and interactions are shown in dashed gray outline; these represent physical interactions that do not correspond to negatively correlated fitness profiles.

Because positive correlations can capture entire complexes, the positively correlated network can identify novel complex members that interact physically and share similar fitness effects (**Figure 7D-I**). Examples include C18orf21, which links physically and functionally to the RNAse P complex (**Figure 7D**); and C19orf25, which links GARP and SNARE complexes (**Figure 7F**). While these are observed in both 293T and HCT116 cells, positively correlated interactions involving protocadherins (**Figure 7E**) and the unknown protein C10orf88 (**Figure 7G**) vary across cell lines.

In contrast, negatively correlated interactions capture specific regulatory relationships. Examples include MDM2 and MDM4, key negative regulators of TP53 (**Figure 7J**); alternating rings of pro- and anti-apoptotic proteins (**Figure 7K**); mixtures of CDK’s, cyclins, and CDK inhibitors (**Figure 7M**); and inhibition of the P-TEFb complex (CDK9, CCNT1, CCNT2) by incorporation into the 7SK snRNP complex (**Figure 7N**). In sum, in combination with BioPlex, Achilles fitness profiles reveal functional relationships among physically interacting proteins that can drive biological discovery.

## DISCUSSION

### The BioPlex Networks Profile Protein Interactions with Unparalleled Depth

Despite considerable effort, our understanding of the interactome is incomplete. By deploying affinity enrichment at scale, our AP-MS platform has profiled interactions at unparalleled depth, encompassing 70% of known human proteins and expanding our knowledge of interactions involving poorly characterized proteins (**Figure 1B-C**). Moreover, through detection in compatible cell contexts, AP-MS can preserve factors such as post-translational modifications and co-complex members that can initiate or stabilize interactions, facilitating detection of complexes (**Figures S1A-C, 3F-I**). This enhances our ability to detect new communities (**Figures 1D, 4**) and to associate new proteins with known complexes (**Figures 1D, 6, S7**).

Because the interactome is dynamic, complete coverage is not achievable in a single experimental context, regardless of methodology. Thus, profiling interactions in another cell type has expanded interactome coverage (**Figure 2**). These additional interactions have revealed many cell-specific pathways and modules, enhancing the scope of the combined BioPlex network (**Figures 3-5**, **7, S2, S5, S6**). Identification of context-specific interactions is especially important, as current knowledge favors core complexes with broadly essential functions (**Figures 3-4**).

### Proteome-scale 293T and HCT116 Networks Reveal Interactome Cell-type Specificity

That protein interactions vary with context is a fundamental biological principle, established through decades of biochemical studies and exemplified by nuclear pore complex customization (Ori et al., 2013). Yet our understanding of interactome dynamics remains rudimentary. While incorporating AP-MS in a second cell line has expanded interactome coverage and validated thousands of interactions, its greatest significance is enabling comparison of cell-line-specific, proteome-scale interaction networks acquired using consistent methodology. Viewing individual proteins (**Figures 2E-H, S2F-I, S6**), subnetworks of shared function (**Figures 3, 4, 5, S5**), and even entire networks (**Figures 2, S3**) reveals widespread reorganization in support of each cell’s unique physiology. Whereas shared interactions reside in dense subnetworks and preferentially link consistently expressed, often essential proteins with long evolutionary histories, cell-specific interactions frequently span disparate network components, coupling variably expressed proteins in context-specific configurations. Interactions among proteins of shared function often vary in tandem, assembling specialized modules according to need. These direct observations of interactome re-organization complement reports of co-variation within protein complexes as inferred from constituent protein expression across cell lines, tissues, and individuals (Ori et al., 2016; Romanov et al., 2019; Ryan et al., 2017). Such extensive context specificity also aligns with disease-specific interactions of the cystic fibrosis transmembrane receptor (Pankow et al., 2015) and with interactome reorganization across tissues (Luck et al., 2019; Skinnider et al., 2018) and upon interferon stimulation (Kerr et al., 2019).

Extensive proteome coverage has been essential to establishing patterns of interactome reorganization. In part this is because large swaths of interaction space reorganize across cell types, especially outside core complexes. More importantly, we have focused our analyses on entire subnetworks rather than individual proteins. While cell-specific differences in an IP might signify either technical or biological variability, tandem variation within GO categories, pathways (**Figures 3, S5**), communities (**Figure 4**), and domain-domain associations (**Figure 5**) across many independent AP-MS experiments suggests a biological basis.

From a systems biology perspective, out networks connect cellular phenotypes to variations in network structure. For example, DREAM complex dissociation reflects SV40 large T antigen expression that drives immortality in 293T cells (**Figure 3C**). Similarly, pre- and post-synaptic membrane complexes are detected in 293T cells (**Figure S5A-B**), evoking their proposed neural crest lineage (Stepanenko and Dmitrenko, 2015). Finally, 293T and HCT116 networks reflect their mesenchymal and epithelial character (**Figures 5G, 7E, S2F, S5E**). If a fundamental goal of proteomics is to biochemically link genotype to phenotype, defining the context-specific interactome is essential.

### The BioPlex Networks Enable Biological Discovery

One motivation for mapping the interactome is that its structure encodes diverse biological insights (**Figure 1D**). By incorporating over 100,000 previously unknown interactions that span 70% of the known proteome (**Figure 1C**), BioPlex suggests function, localization, and complex membership for thousands of proteins, such as CASTOR1/2 (Chantranupong et al., 2016) and SAMTOR (Gu et al., 2017). In fact, as BioPlex has grown, its utility for biological discovery has increased via two mechanisms. Most obviously, additional baits increase coverage and enhance a network’s predictive power as noted previously for disease gene discovery (Huang et al., 2018). More subtly, as BioPlex has grown the same complexes have been immuno-purified multiple times, affirming associations through repeated co-purification (**Figure S1G-I**). Such intra-network confirmation now supports more than half of all BioPlex interactions. These internally validated interactions are especially attractive for biological follow-up, as shown for C11orf49 and the TTLL1-polyglutamylase complex (**Figures S1G, 6, S7**).

While BioPlex 3.0 enables biological discovery, the HCT116 network enhances its scope and predictive power. First, by incorporating additional interactions and proteins, the HCT116 network enhances proteome coverage. This is especially important since interactions among entire functional classes may be absent in individual cell lines. Second, over 35,000 interactions are confirmed in a second cell line and thus more likely to be authentic and relevant to a complex’s core function. A third way to enhance the predictive power of BioPlex is to combine predictions across networks, as demonstrated by identifying accessory proteins for protein complexes (**Figure 6**). Finally, combining BioPlex networks with cellular fitness data identifies interactions defined by both physical and functional associations (**Figure 7**).

### Toward a Broader View of Interactome Dynamics

Our platform has enabled us to target half of human proteins for AP-MS and to extend this analysis into a second cell line, greatly increasing interactome coverage and enabling us to compare multiple proteome-scale, context-specific interaction networks. Despite these advances, our understanding of the interactome remains incomplete. The first limitation is protein coverage: while we have completed AP-MS experiments for over 10,000 human proteins, nearly as many remain untargeted. Ongoing development of human gene libraries promises to make these remaining proteins accessible, though their AP-MS analysis remains onerous.

More fundamental are technical limitations of AP-MS. Some protein classes, especially membrane proteins, are difficult to purify and the affinity tag itself may introduce artifacts. While specialized protocols could be used for membrane proteins and tag effects could be resolved through AP-MS of N-terminally tagged proteins, both are challenging at proteome scale. Another limitation is that affinity purification precludes detection of low-affinity binding partners. Proximity labeling could capture weaker, more transient interactions and aid study of multi-pass membrane proteins. Indeed, large-scale proximity labeling is underway to define proteome spatial organization (Go et al., 2019), providing a complementary view of protein interactions.

While scale and technical limitations are significant, most challenging are interactome dynamics. Here we have shown that interactomes vary with cellular context. Yet no two cell lines can capture the proteome diversity observed across hundreds of cell types, stimuli, and growth conditions. Since each cell expresses roughly half of all human proteins, it can only cover a fraction of the interaction space. Systematic AP-MS will be essential for probing context-specific interactomes with maximum depth. However, because AP-MS is time- and labor-intensive, parallel deployment of complementary experimental approaches like co-fractionation (Heusel et al., 2019) will maximize interactome coverage.

We have reported the first AP-MS analysis of half the human proteome, generating two proteome-scale, cell-specific interaction networks. This has broadened our understanding of the protein interaction landscape and defined how the interactome adapts to variable cellular contexts. While this work just begins to explore cell specificity of the interactome, it demonstrates the importance of interaction profiling across cellular contexts and establishes a foundation for analysis of interactome dynamics.

## AUTHOR CONTRIBUTIONS

This study was conceived by S.P.G. and J.W.H. E.L.H. developed *CompPASS-Plus* and software for data collection and integration, performed all bioinformatic and statistical analyses and oversaw data collection and AP-MS pipeline quality. R.J.B. and L.P.V. managed the cell culture and biochemistry. J.N.-P., J.C., S.G.T. and J.A.P. were responsible for mass spectrometry operation. K.B., F.G., M.P.G., H.P., J.S., S.T. A.T., S.F., A.G., E.M., K.S. and A.P. performed cell culture. R.J.B., G.Z., K.B. and E.M. performed all affinity purifications and prepared samples for MS analysis. D.K.S. developed BioPlexExplorer. R.R. provided computational support while J.P. and D.P.N. assisted with bioinformatic analyses. A.T. and L.P.V. performed validation experiments. The paper was written by E.L.H., J.W.H. and S.P.G. and was edited by all authors.

## ACKNOWLEDGEMENTS

We thank Marc Vidal and David Hill for ORFeome 8.1 and acknowledge the Nikon Imaging Center (Harvard Medical School) for imaging support. This project was supported by the NIH (U24 HG006673 to S.P.G., J.W.H., and E.L.H.) and Biogen (S.P.G. and J.W.H.).

## STAR METHODS

### LEAD CONTACT AND MATERIALS AVAILABILITY

Further information and requests for resources and reagents should be directed to and will be fulfilled by the Lead Contact, Steven Gygi.

### EXPERIMENTAL MODEL AND SUBJECT DETAILS

#### Clone Construction

Clones corresponding to human genes were obtained and prepared for AP-MS analysis as described previously (Huttlin et al., 2015). Briefly, open reading frames were taken from the human ORFeome, version 8.1 (Yang et al., 2011) and the OC collection (http://horfdb.dfci.harvard.edu) and cloned using Gateway techniques into a lentiviral recipient vector that incorporated a c-terminal FLAG-HA tag and expressed the target ORF and a puromycin resistance marker under control of a CMV promoter. All clones were sequenced to verify ORF identity.

Within version 8.1 of the ORFeome, clones were generally sorted by length and accordingly assigned to 96-well-plates. In contrast, OC collection baits were arrayed on 96-well plates in more random order. Throughout the project, clones have generally been targeted for AP-MS in batches corresponding to these 96-well plates. During initial AP-MS analysis in 293T cells, plates were run in random order, though plates corresponding to very large or small clones were avoided until late in the project. For HCT116 cells, roughly half of plates consisted of cherry-picked baits that 1) had worked in 293T cells and 2) were selected to sample evenly across the 293T interaction network; subsequently we have run remaining baits in batches according to their original ORFeome plate assignments. Baits repeated in 293T cells were also selected to sample evenly across the 293T interaction network.

#### Creation of Stable Cell Lines in 293T and HCT116 Cells

Cell culture and bait expression were performed as described previously (Huttlin et al., 2015). Briefly, lentivirus was produced in 293T cells following transfection of Gateway expression clones using PEI as an adjuvant. Upon subsequent lentiviral transduction of bait constructs, either 293T cells or HCT116 cells (both from ATCC) were expanded in the presence of 1 µg/mL puromycin selection (Gibco) to obtain five 10-cm dishes per cell line. Once they reached minimum 80% confluence, cells were washed with ice cold PBS phosphate-buffered saline (PBS) pH 7.2 (Gibco) and harvested via gentle scraping, followed by centrifugation to pellet cells and PBS removal. Cell pellets were stored at −80°C.

All cell lines were tested for presence of mycoplasma using the Mycoplasma Plus PCR assay kit (Agilent) and found to be free of contamination. Additionally, the identities of 293T and HCT116 cell lines were verified through GTG-banded karyotyping by the Brigham and Women’s Hospital Cytogenomics Core Laboratory.

## METHOD DETAILS

### Affinity-Purification Mass Spectrometry

Affinity-purification mass spectrometry experiments were performed as previously described in detail (Huttlin et al., 2015, 2017). Methods are summarized below. AP-MS experiments in each cell line were performed separately: HCT116 AP-MS experiments were not initiated until all experiments in 293T cells had been completed. Baits targeted in 293T cells are summarized in **Table S1A** while baits targeted in HCT116 cells are summarized in **Table S2B**.

### Affinity Purification of Protein Complexes

After thawing, cell pellets were lysed in 50 mM Tris-HCl pH 7.5, 300 mM NaCl, 0.5% (v/v) NP40 and the lysate was clarified by centrifugation and filtration. Immobilized, pre-washed mouse monoclonal anti-HA agarose resin (Sigma-Aldrich, clone HA-7) was used to immunoprecipitate affinity-tagged bait proteins and their binding partners. Beads were incubated with lysates for 4 hours at 4°C, followed by supernatant removal, four washes with lysis buffer, and two washes with PBS (pH 7.2). Elution was achieved in two steps via addition of 250ug/mL HA peptide in PBS at 37°C for 30-minute incubations with gentle shaking.

Following affinity enrichment, residual HA peptide and other non-protein material were removed through precipitation with 20% TCA followed by a wash with 10% TCA and triplicate washes with cold acetone. Pellets were then resuspended and reduced in 150 µL of 100 mM ammonium bicarbonate (Sigma-Aldrich) with 1 mM DTT (Fluka) and 200 ng of sequencing grade trypsin (Promega) was added for digestion overnight at 37°C. For HCT116, TCA precipitated pellets were resuspended in 20uL of 55mM Tris pH 8.5/10% Acetonitrile/1mM DTT, with 200ng trypsin added per sample. 5% HPLC-grade formic acid (Thermo Fisher Scientific) was subsequently added to quench digestion and samples were desalted with homemade stage tips as described previously (Rappsilber et al., 2007). After elution with 80% acetonitrile/5% formic acid, peptides were dried in a speed-vac and resuspended in 16µL of 5% formic acid and 4% acetonitrile.

### Mass Spectrometry Data Acquisition

All mass spectrometry data were acquired on first-generation Q-Exactive mass spectrometers (Thermo Fisher Scientific) equipped with Famos autosamplers (LC Packings) and Accela600 liquid chromatography (LC) pumps (Thermo Fisher Scientific). Microcapillary columns used for peptide separation were prepared in-house by packing ∼0.25 cm of Magic C4 resin (5 µm, 100 Å, Michrom Bioresources) and ∼18 cm of Accucore C18 resin (2.6 µm, 150 Å, Thermo Fisher Scientific) into a 100 µm inner diameter microcapillary column. We loaded 4 µL of sample onto the column for each analysis. Each sample was run in technical duplicate in accordance with the requirements of the *CompPASS-Plus* algorithm (described below).

Each LC-MS run was approximately 70 minutes long including sample loading, analytical separation, and column re-equilibration. Peptide separation occurred over 40 minutes using a gradient from 5 to 26% acetonitrile in 0.125% formic acid. The scan sequence consisted of a single MS1 spectrum followed by MS2 scans targeting up to twenty precursors. Orbitrap MS1 scans were acquired in centroid mode at 70,000 resolution spanning a range from 300-1500 Th with the automatic gain control (AGC) target set to 1.0e6 and a maximum ion injection time of 250 ms. Following higher-energy collision-induced dissociation (HCD), MS2 spectra were acquired in centroid mode at 17,500 resolution with the AGC set to 1.0e5 and a maximum ion injection time of 60 ms. The isolation window was set to 2 Th and the normalized collision energy (NCE) was 25-30. Features were targeted for MS2 analysis in order of decreasing intensity while excluding unassigned and singly charged features. Dynamic exclusion was set to automatic.

### Identification of Peptides and Proteins

Upon completion of each LS-MS analysis, the resulting RAW files were converted to mzXML format using msconvert (ProteoWizard). Monoisotopic peak assignments and charge states were then verified for all features targeted for MS2 analysis and MS/MS spectra were matched with peptide sequences using the Sequest algorithm (Eng et al., 1994) along with a composite sequence database including the human Uniprot database (Consortium, 2015), GFP (our negative control), the FLAG-HA affinity tag, and sequences of common contaminants. This Uniprot database includes both SwissProt and Trembl entries and dates to the outset of this AP-MS study in 2013. Protein sequences were listed in both forward and reversed order to facilitate false discovery rate control. During database searching, fully tryptic peptides with up to two missed cleavages were considered and MS1 and MS2 tolerances were set to 50 ppm and 0.05 Da, respectively. Variable oxidation of methionine (+15.9949) was permitted.

Following Sequest analysis, peptides and proteins identified in each LC-MS analysis were filtered in two steps using the target-decoy method (Elias and Gygi, 2007) to control both peptide- and protein-level false discovery rates. First, a linear discriminant function was trained to distinguish correct and incorrect peptide identifications using Xcorr, DCn, peptide length, charge state, mass error, missed cleavage count, and fraction of ions matched (Huttlin et al., 2010) and peptides were filtered to a 1% FDR. Next, these peptides were assembled into proteins, scored probabilistically, and filtered to a 1% protein-level FDR.

Although the filters previously described controlled the peptide- and protein-level FDR within each individual LC-MS analysis, when many datasets are combined as described in this study, the overall dataset-level FDR tends to balloon as false positives accumulate. Since our 293T and HCT116 datasets include over 20,000 and 10,000 LC-MS runs respectively, without additional filtering this problem would be especially severe. To further control the global protein FDR as AP-MS datasets were combined, entropy-based filtering was applied to remove proteins inconsistently detected across technical replicates (Huttlin et al., 2015) and protein identifications within each IP were required to be supported by multiple unique peptides when technical replicates were combined. Together these additional filters reduced the global FDR by more than 100-fold.

### AP-MS Pipeline Quality Control

Several steps have been taken to ensure that sample preparation and instrument performance has been maintained at consistent levels throughout the duration of this project. First, standards were routinely run on the instrument to monitor instrument performance over time. These standards consisted of trypsin-digested yeast whole cell lysate that was diluted to approximate the complexity and sample load of a typical AP-MS sample. Second, positive and negative control IP’s (RAB11B and GFP) were analyzed with every plate. Third, every IP was analyzed in technical duplicate with replicate runs acquired in reverse order on different LC-MS systems. These replicate analyses were primarily done to aid LC carry-over removal and enhance detection of high confidence interacting proteins, though they also enabled us to continuously compare the relative performance of all instruments devoted to this project in real time, making detection of instrument problems much easier.

In addition, every LC-MS run acquired for this project was screened using an automated anomaly detection system to identify problematic runs following database searching as described below. This algorithm monitored a variety of statistics for each run such as numbers of PSM’s, unique peptides, and proteins identified, estimated maximum numbers of true positives (TPMax), missed cleavage rate, average Xcorr, numbers of MS2 spectra acquired, etc. Each run was compared with all prior runs with respect to each of these parameters and scored using a kernel-density-based probabilistic model; anomalous runs were filtered allowing 1% of runs flagged as anomalies to be false positives. This algorithm updated continually as runs were added, allowing the algorithm to adapt to changing conditions. Classifications were monitored and manually corrected as necessary to maintain accurate performance. To accommodate cell-line-specific differences, separate models were trained from runs corresponding to 293T cells and HCT116 cells.

Prior to *CompPASS-Plus* analysis, each IP was screened to ensure detection of the expected bait protein, thus protecting against expression problems and occasional sample mix-ups. We expected each bait to be detected at levels significantly higher than background when targeted for AP-MS enrichment. Thus, we required the expected bait to be detected at elevated levels in two technical replicates for an AP-MS experiment to be eligible for *CompPASS-Plus* scoring. For a small number of baits whose sequences do not directly correspond to Uniprot entries, custom searches were performed to verify bait identities. These additional searches were only used for bait verification and only searches against the standard Uniprot database were used for interaction network generation. If the expected bait was not confidently detected and could not be determined to result from a mix-up, the runs were discarded, and the IP was repeated.

Before an AP-MS experiment could be included in CompPASS-Plus analysis, two technical replicate LC-MS analyses passing all quality control checks were required. To accommodate occasional low-quality runs, we scaled our AP-MS pipeline to offer quadruplicate analysis. This provided up to two additional runs in cases where poor instrument performance compromised one of the initial runs. If two acceptable runs were not obtained after up to four injections, the bait was targeted for repeat AP-MS analysis.

### Inferring Protein Interactions from LC-MS Data

Here we provide an overview of our approaches for identifying high-confidence interacting proteins from AP-MS data and creating integrated interaction networks as published previously (Huttlin et al., 2015). These procedures were performed separately for AP-MS data acquired in 293T and HCT116 cells. Additional information is provided below regarding how these networks were combined for comparative analysis. The BioPlex 3.0 network results from re-analysis of all AP-MS data in BioPlex 1.0 and 2.0, along with 4,237 additional new AP-MS datasets.

#### Identification of High Confidence Interacting Proteins

Identification of high confidence interacting proteins proceeded in four steps: 1) merging technical replicates; 2) CompPASS analysis; 3) Post-CompPASS filtering; and 4) CompPASS-Plus analysis. First, those AP-MS experiments that passed all quality control filters were identified; for each, their associated technical replicates were combined to produce a summary of proteins identified. Across replicates, peptides were re-assembled into proteins according to principles of parsimony and Uniprot ID’s were mapped to Entrez Gene ID’s. All subsequent analysis was performed on data assembled at the Gene ID level to address complications due to protein isoforms. For each gene product, spectral counts were averaged across replicates and entropy scores were calculated as described previously (Huttlin et al., 2015).

Second, all AP-MS experiments were scored using *CompPASS* essentially as described previously (Behrends et al., 2010; Sowa et al., 2009). Merged data from all AP-MS experiments matching a single cell line were combined to create a “stats” table that indicates the number of PSM’s observed for each gene product in each AP-MS experiment. *CompPASS* analysis produced two scores for each protein detected in each IP: a z-score that reflects a given protein’s abundance compared to its typical background levels across all other IP’s in the dataset; and an NWD score, which takes each gene product’s abundance, detection frequency, and technical replication into account to estimate its enrichment compared to all other IP’s. NWD scores were scaled to assign values of 1.0 or higher to the top 2% of candidate interacting partners.

Third, after *CompPASS* analysis was complete, additional filters were applied to protein identifications in each IP to remove inconsistent or low-confidence protein identifications and avoid false positive interactions. These filters, two of which are also described above in the section titled “Identification of Peptides and Proteins” include 1) discard proteins for which only a single unique peptide sequence was observed across two technical replicates; 2) require a minimum entropy score of 0.75 (calculated using Log2) comparing spectral counts observed in two technical replicates; and 3) within each 96-well-plate, look for proteins detected with unusual frequency compared to a background defined by all other plates – if statistically significant enrichment is detected, discard observations on that plate that fall below its average. The first two of these filters help to control the global protein-level false discovery rate; furthermore, filters 2 and 3 protect against LC carry-over, especially as observed for overexpressed baits; finally, filter 3 protects miscellaneous plate-specific variations in protein detection.

Fourth, following *CompPASS* scoring and application of filters as described above, all remaining bait-prey associations from a single cell line were scored and classified using the supervised classifier *CompPASS-Plus* as described previously (Huttlin et al., 2015). To enable training, each bait-prey association was assigned one of three preliminary labels (false positive interaction, background protein, or specific interactor). All preys corresponding to decoy protein sequences were labeled as false positives; preys whose interactions were confirmed in STRING (Szklarczyk et al., 2017) or GeneMANIA (Franz et al., 2018) were labeled as true positives; all others were labeled as background. In addition, because their levels are strongly enriched by design following affinity purification, baits within each IP were also labeled as “true interactors” for modeling purposes. Moreover, because both published and unpublished versions of BioPlex data have been previously released, care was taken to ensure that the STRING and GeneMANIA data used for training did not incorporate the results of any prior BioPlex analyses. Specifically, we used archival versions of these databases that predated release of any BioPlex datasets.

Once these preliminary labels were assigned, a separate Naïve Bayes classifier was trained for each 96-well plate matching either 293T or HCT116 cells, using all other 96-well-plates from that cell line for training. This plate-based, leave-one-out cross-validation scheme ensured that separate data were always used for model training and scoring. Features considered by the classifier included *CompPASS* NWD and Z-scores, entropy, a plate-based z-score, binned unique peptide counts, the fraction of IP’s in which a given protein was detected, the total number of PSM’s observed for a given protein across all IP’s, the ratio of observed PSMs in a given IP to the total number of PSM’s across all IPs, and the unique:total peptide ratio within the given IP. The output of this algorithm was a vector of three scores reflecting the likelihood that each interaction resulted from either an incorrect protein identification, background, or a *bona fide* interacting partner. *CompPASS-Plus* was implemented in R (RCoreTeam, 2011) using the Naïve Bayes classifier in the package e1071 (Meyer et al., 2012). Because many features did not conform to normal distributions, continuous features were discretized by splitting each into 1000 equally-sized bins for classification.

#### Interaction Network Assembly

Following *CompPASS-Plus* scoring, all AP-MS experiments corresponding to a single cell line were assembled into a network that models the human interactome. While previous versions of BioPlex excluded a small number of baits with very high numbers of interacting partners (> 100), no filter was applied limiting the numbers of interactions that could be identified in a single AP-MS experiment. In cases where the same bait protein had been targeted multiple times, often because the ORFeome contained more than one matching clone, only a single IP was retained. In these cases we preferentially kept IP’s corresponding to full-length clones and favored IP’s that returned more interacting partners. Individual interacting partners were also filtered as the network was assembled. Specifically, a few dozen remaining decoy proteins were removed, as were common contaminant proteins (e.g. keratin).

Once filtering was complete, a nonredundant list of observed bait-prey pairs was compiled accompanied by *CompPASS-Plus* scores for each. When pairs of baits were found to associate reciprocally, their *CompPASS-Plus* scores were merged as described previously (Huttlin et al., 2015) to increase the probability of interaction. Finally, this list of bait-prey pairs was filtered according to *CompPASS-Plus* score: interactions with an interaction score equal to or greater than 0.75 were retained to create the final interaction network. This cutoff of 0.75 corresponds to the top ∼2% of candidate interactions in both 293T and HCT116 cells. Interactions observed in 293T and HCT116 networks are summarized in **Table S1B** and **Table S2C**, respectively.

### Assembly of Combined and Replicated Networks

As described above, 293T and HCT116 networks were initially assembled independently through separate *CompPASS* and *CompPASS-Plus* analyses. To facilitate comparison, these networks were subsequently aligned using the following procedure:

1. Interactions within both 293T and HCT116 networks were merged to generate a single non-redundant list. For this purpose, each interaction was represented as a pair of linked Entrez Gene ID’s. This step effectively defines the “Combined Network” shown in **Figure 2D** and includes every interaction among the top 2% detected in 293T cells, HCT116 cells, or both.
2. Each interaction in the “Combined Network” was looked up in both 293T and HCT116 datasets. By definition, each of these interactions scored in the top 2% for at least one cell line, as was required for inclusion in the original networks. If this interaction ranked among the top 2% in 293T cells and was also observed among the top 5% of candidate interactions in HCT116 cells, then it was counted as replicated in both cell lines. Similarly, if the interaction ranked among the top 2% in HCT116 cells and was also observed in the top 5% of 293T cells, it was counted as replicated. Together these define the “shared interactions” and the “Replicated Network” in **Figure 2D**.
3. If the interaction ranked among the top 2% of candidate interactions in one cell line and was either undetected or failed to score among the top 5% of candidate interactions in the other cell line, then it was considered either 293T- or HCT116-specific as shown in **Figure 2D**.
4. a slightly relaxed threshold for judging whether edges were replicated in the opposite network was necessary due to the inherent challenges of identifying a relatively small number of *bona fide* interacting partners within a much larger set of background interactions. This relaxed threshold for replication enabled us to be confident that edges we call cell-line specific correspond to substantial differences in detection rather than ‘near-misses’ that were detected and fell just short of our chosen significance cutoffs.

A complete list of interactions within this combined network is provided in **Table S2C**; each edge is labeled to indicate whether it maps to the original 293T or HCT116 networks and whether it was classified as replicated or cell-line-specific as described above.

#### 293T Replication Experiments

Baits selected for replicate AP-MS analysis in 293T cells were selected from the set of baits that worked following first-pass analysis in 293T cells (**Table S2E**). Baits were selected to sample evenly across the interactome, ensuring that large complexes such as the ribosome and proteasome would not be over-represented in the replication set and that baits falling outside of well-defined clusters would be included as well. Identical virus was used to infect a second batch of 293T cells. Cell culture, affinity purification, and mass spectrometric analysis were all performed using standard procedures as described above. These replicates were run on separate 96-well plates separated by several months to several years from the original AP-MS analysis. Replicate datasets were scored against the main 293T “stats” table just as the original IP’s were, though they were not included in the stats table for scoring other IP’s. Identical filters for unique peptides, entropy scores, etc. were applied to replicates. And replicate IP’s were scored using *CompPASS-Plus* models trained on the full 293T dataset using plate-based leave-one-out cross-validation as described previously. For consistency with our 293T – HCT116 comparisons, interactions identified in the original 293T IP’s were considered replicated if they were detected in the top 5% in the second IP (**Table S2F**).

### Quantitative Proteomic Comparison of 293T and HCT116 Cells

The quantitative proteomic comparison of 293T and HCT116 proteomes presented here has been published previously (Erickson et al., 2019). Protein extraction, digestion, and TMT labeling were accomplished using the SL-TMT method (Navarrete-Perea et al., 2018). First, quintuplicate pellets of 293T and HCT116 cells were lysed in 8M urea and lysates were reduced with 5 mM TCEP and alkylated with 10 mM iodoacetamide. Proteins were then purified via methanol-chloroform precipitation, resuspended in 200 mM EPPS, pH 8.5 and digested sequentially with LysC and Trypsin. Peptides were then labeled with TMT, combined, and desalted prior to high-pH reversed-phase fractionation to produce 24 fractions. Of these, 12 non-adjacent samples were analyzed via LC-MS.

All samples were analyzed on an Orbitrap Fusion Lumos utilizing an MS3 workflow that featured a custom real-time database search method as described previously (Erickson et al., 2019). Briefly, the instrument was operated in data-dependent mode with each MS1 scan followed by multiple MS2 scans to attain a cycle time of 2 seconds. MS1 spectra were collected in the Orbitrap at 120,000 resolution allowing ion times up to 100 ms. Features with defined monoisotopic masses and charge states greater than one were targeted for collision-induced dissociation in descending order based on intensity. MS2 spectra were acquired in the ion trap using a quadrupole isolation width of 0.5 m/z; automatic gain control of 20,000; and maximum ion time of 35 ms.

Upon acquisition, each MS2 spectrum was provided via API to a custom module that performed database searches in real time. Each spectrum was searched against a database containing Uniprot human sequences plus common contaminants. When a spectrum matched a human peptide sequence with binomial score of at least 55, an MS3 scan was triggered through the API. MS3 spectra were acquired for up to 150 ms at an AGC of 50,000 utilizing a normalized collision energy of 55. Synchronous precursor selection of up to 10 fragments was employed; all selected fragments were required to account for at least 5% of base peak signal and to match a b- or y-ion corresponding to the predicted peptide.

Following acquisition, RAW files were converted to mzXML format using msconvert. Monoisotopic peak assignments were then verified for any features that were targeted for MS2 analysis. Spectra were then submitted for database searching using SEQUEST (Eng et al., 1994) along with a database of human protein sequences (Uniprot, 2014) and common contaminants in forward and reversed orientation. Fully tryptic peptides were considered for database searching assuming precursor and product ion tolerances of 50 ppm and 0.9 Th, respectively. Fixed modifications of TMT (+229.163 Da) on lysines and N-termini and carbamidomethylation (+57.021 Da) of cysteines were assumed while variable oxidation (+15.995 Da) of methionine was considered. Peptide-spectral matches were filtered to a 1% FDR using the target-decoy method in combination with linear discriminant analysis (Huttlin et al., 2010); subsequently, peptides were assembled into proteins and filtered to attain a 1% protein-level FDR and to account for peptide redundancy according to principles of parsimony. TMT quantitation of each protein was performed by summing reporter ion intensities across all matching peptide-spectral-matches with corresponding TMT data; only peptides that attained a minimum summed reporter ion signal-to-noise of at least 100 were retained for quantitation. TMT channels were subsequently normalized assuming equal protein loading. Finally, each protein’s TMT profile was scaled to sum to 100%, thus reporting the fractional TMT signal associated with each channel. The final dataset is provided in **Table S2A**.

### C11orf49 Validation Experiments

#### Affinity Purification and Immunoblot analysis

293T or HCT116 cells were transduced with c-terminal FLAG-HA lentiviral constructs harboring previously characterized cDNAs of the TTLL1 polyglutamylase complex or C11orf49, a candidate complex member identified in this study. Cells were selected with puromycin as described above, expanded to five 10cm plates, and harvested via gentle scraping into PBS. Frozen cell pellets were lysed with 50mM Tris pH 7.5, 150mM NaCl, 1% NP-40, 1mM DTT), with protease inhibitors (Roche, Complete mini EDTA free) on ice. Lysates were cleared by centrifugation, and subjected to affinity purification using either HA-agarose (Sigma) or anti-FLAG magnetic beads (Sigma) for 2 hours at 4°C. Beads were washed 4x with lysis buffer, then complexes were eluted with 3xFLAG peptide or beads were boiled in 1x NuPage sample buffer (Invitrogen) prior to SDS-PAGE. Proteins were transferred to PVDF membranes and immunoblotted with the following antibodies: C11orf49 (Cat. No. 20195-1-AP, Proteintech), TPGS1 (ab184178, Abcam), and anti-FLAG (M2)-HRP (A8592, Sigma).

#### Confocal Microscopy

U2OS cells (American Type Culture Collection) were plated on glass coverslips (Zeiss) and transduced with lentiviral vectors expressing C-Flag–HA-tagged baits. At 48 h after infection, cells were fixed with 4% paraformaldehyde for 15 min at room temperature. Cells were washed in PBS, then blocked for 1h with 5% normal goat serum (Cell Signaling Technology) in PBS containing 0.3% Triton X-100 (Sigma). Coverslips were incubated with anti-HA antibodies (mouse monoclonal, clone HA.11, BioLegend) or anti-HA plus anti-PCM1 (#5213, Cell signaling) for 2 h at room temperature. Anti-C11orf49 (20195-1-AP, Proteintech) and anti-γ tubulin (ab11317, Abcam) were utilized to visualize C11orf49 localization in proximity to centrosomes following coverslip permeabilization in ice cold methanol for 20 minutes. Cells were washed three times with PBS, then incubated for 1 h with appropriate Alexa Fluor-conjugated secondary antibodies (ThermoFisher). Nuclei were stained with Hoechst, and cells were washed three times with PBS and mounted on slides using Prolong Gold mounting media (ThermoFisher). All images were collected with a Yokogawa CSU-X1 spinning disk confocal with Spectral Applied Research Aurora Borealis modification on a Nikon Ti-E inverted microscope equipped with a Nikon 100 × Plan Apo numerical aperture 1.4 objective lens (Nikon Imaging Center, Harvard Medical School). Both confocal and widefield images were acquired with a Hamamatsu ORCA-ER cooled CCD (charge-coupled device) camera controlled with MetaMorph 7 software (Molecular Devices). Fluorophores were excited using a Spectral Applied Research LMM-5 laser merge module with acousto-optic tuneable filter (AOTF)-controlled solid-state lasers (488 nm and 561 nm). A Lumencor SOLA fluorescence light source was used for imaging Hoechst staining. *z* series optical sections were collected with a step size of 0.2 μm, using the internal Nikon Ti-E focus motor, and stacked using FIJI (Image J) to construct maximum intensity projections. Image brightness and contrast were adjusted for each image equally among staining conditions using FIJI software.

### QUANTIFICATION AND STATISTICAL ANALYSIS

Unless otherwise indicated, all network analyses were performed in *Mathematica 11.3* or *12.0* (Wolfram Research). All statistical tests have been adjusted for multiple testing correction using the Benjamini-Hochberg method (Benjamini and Hochberg, 1995). All figures were plotted in *Mathematica* and assembled in *Adobe Illustrator*.

### BioPlex 3.0 Analysis (Figure 1)

#### Comparison with BioGRID and PubMed

To determine the extent to which BioPlex 3.0 interactions had been reported previously in the literature, we compared our interaction network against interactions compiled by BioGRID (Oughtred et al., 2018). Human interactions reported in BioGRID version 3.5.167 (November 2018) were downloaded and filtered to remove interactions from previous published and unpublished BioPlex networks. We then searched for each BioPlex interaction in the filtered BioGRID dataset; if a match was found, we next queried whether the interaction was reported in high-throughput or low-throughput studies as recorded by BioGRID. If at least one reference pointed to a low-throughput study, that edge was counted as reported in low-throughput; alternatively, if all references pointed to high-throughput studies, the edge was counted as confirmed through high-throughput studies; finally, if no matches were found independent of previous BioPlex datasets, the edge was counted as unique to BioPlex.

PubMed publication counts associated with each human gene were obtained via FTP from NCBI in December 2018. Proteins in the BioPlex 3.0 network were then assigned to 100 bins corresponding to percentiles based on their associated citation counts. Each interaction in BioPlex was then categorized based on whether it was confirmed in low- or high-throughput or novel to BioPlex and binned according to the percentiles of its two associated proteins, producing the density plots displayed.

#### Functional and Disease Associations

To assess functional and disease associations of proteins in the BioPlex network, we superimposed GO functional data (Ashburner et al., 2000) and DisGeNET disease associations (Piñero et al., 2017). GO assignments were downloaded from www.geneontology.org in November 2018 while DisGeNET disease associations were downloaded from www.disgenet.org in May 2019. For each protein in the BioPlex 3.0 network GO terms or diseases associated with two or more of their neighbors were identified. The enrichment of each term was evaluated relative to its background abundance in the entire network using a one-sided hypergeometric test. Multiple testing correction via the method of Benjamini and Hochberg (Benjamini and Hochberg, 1995) was then applied and results were filtered to a 1% FDR. GO Cellular Components, Molecular Function, and Biological Process terms and DisGeNET disease associations were processed separately. Results are summarized in **Table S1C-I**.

#### Prediction of Subcellular Localization

Localization predictions were performed as described previously (Huttlin et al., 2015). Briefly, subcellular localization data for all human proteins were downloaded from Uniprot (January 2019) and mapped to the BioPlex network after manually merging localization information into thirteen core categories: nucleus, cytoplasm, cytoskeleton, endosome, ER, extracellular, Golgi, lysosome, mitochondria, peroxisome, plasma membrane, vesicle, and cell projection. Subcellular localization predictions were based on the enrichment of each category among a proteins’ primary and secondary neighbors as quantified via Fisher’s Exact Test using contingency table with 2 rows to indicate membership in a particular localization category (yes/no) and 3 columns corresponding to a proteins’ primary neighbors, secondary neighbors, or all other proteins in the network. P-values were adjusted for multiple testing (Benjamini and Hochberg, 1995) and to accept a predicted localization as significant, the relevant localization also had to be observed at rates greater than random chance when primary and secondary neighbors (if any) were tallied separately. Predictions are provided in **Table S1J**.

#### Data-driven Discovery of Protein Communities

As described previously (Huttlin et al., 2017), BioPlex 3.0 was partitioned into communities using Markov Clustering (Enright et al., 2002) with an inflation parameter of 2.15 and the force-connected option set to ‘yes’. Only communities containing 3 or more proteins were retained. BioPlex 3.0 communities are summarized in **Table S1K-L**.

#### Association of New Proteins with Known Complexes

To identify new proteins associated with known complexes, non-redundant CORUM complexes (Giurgiu et al., 2018) downloaded in May 2019 were mapped individually to BioPlex. Each protein linked to one or more known complex members was considered a putative associated protein and was scored for edge enrichment using a hypergeometric model with multiple testing correction as depicted in **Figure 5** and described in the section titled “Assignment of Accessory Proteins to Known Protein Complexes” below. Note that whereas Figure 5 and the description below make predictions by combining 293T and HCT116 networks, here the BioPlex 3.0 network was used alone.

#### Discovery of Associations among Protein Domains

The BioPlex 3.0 network was mined for associations among protein domains as described previously (Huttlin et al., 2015). Briefly, PFAM domains were associated with individual proteins in the BioPlex network according to information obtained from Uniprot in January 2019. All domain pairs connected by two or more interactions were identified and the significance of each association was calculated via Fisher’s Exact Test after incorporating numbers of interactions involving each domain individually and the total number of interactions in the network. The Benjamini-Hochberg method was subsequently applied for multiple testing correction and domain associations were deemed significant at a 1% FDR. Domain associations are summarized in **Table S1M**.

#### Network Structural Analysis

Analyses of BioPlex 3.0 network structure were performed in Mathematica 12.0. The vertex degree distribution was fit to a power law via maximum likelihood as described previously (Huttlin et al., 2015). Vertex separation statistics were calculated from pairwise distances among all nodes in the BioPlex 3.0 network. Centrality analysis of essential proteins was performed as follows. First, eigenvector centralities were calculated for each node and proteins were sorted accordingly. Next, broadly essential human genes were identified from fitness effects associated with their CRISPR knockout or RNAi knockdown across hundreds of cancer cell lines as measured in the Achilles project (Tsherniak et al., 2017). Specifically, “common essential” genes identified previously (Dempster et al., 2019) were used to separate the ranked centrality list into two distributions corresponding to “essential” and “non-essential” proteins and these distributions were compared using a Kolmogorov-Smirnov test. The 1,882 “essential” proteins identified in the BioPlex network showed a strong propensity to occupy the most central positions in the network.

### BioPlex 3.0 Validation (Figure S1)

#### Interaction Enrichment among CORUM Complexes

CORUM (Giurgiu et al., 2018) is a manually curated database that reports core protein complexes that is frequently used as a gold standard. Each complex is reported as a list of proteins that assemble to form each complex. We employed a statistical approach to measure the extent to which each CORUM complex was reflected in the architecture of BioPlex and other human interaction networks. The basic assumption of our analysis is that the proteins within a CORUM complex will interact with each other with a frequency that significantly exceeds global connectivity of the network.

For this analysis, each human complex in CORUM’s core complex list (coreComplexes.txt; downloaded in September 2018) was mapped to the interaction network individually. To be eligible for scoring, CORUM complexes were required to have at least 3 members, one or more of which were targeted as baits in BioPlex 3.0. For each target complex, the subnetwork defined by its members was extracted and the interactions within were counted. The enrichment of interactions among target complex members was calculated using a one-sided binomial test assuming a background probability of interaction equal to the interaction density of the interaction network. This analysis returned p-values that were adjusted for multiple testing (Benjamini and Hochberg, 1995). A complex was considered enriched with interactions when the p-value was less than 0.01 after multiple testing correction.

In addition to BioPlex 3.0, we used this method to assess CORUM complex enrichment in several published datasets, including BioPlex 1.0 (Huttlin et al., 2015) and BioPlex 2.0 (Huttlin et al., 2017), along with other datasets acquired via AP-MS (Hein et al., 2015), yeast-two-hybrid (Luck et al., 2019; Rolland et al., 2014; Rual et al., 2005), and correlation profiling (Havugimana et al., 2012; Wan et al., 2015). In addition, we compared scored HuMAP (Drew et al., 2017), which was derived from re-analysis of data combined from several of these other studies (Hein et al., 2015; Huttlin et al., 2015; Wan et al., 2015). Importantly, to enable comparisons across datasets, identical complexes were scored in each. This means sometimes complexes were scored in a relatively small network even when its constituent proteins were not detected or when none of its member proteins had been targeted for AP-MS analysis. The CORUM coverages reported thus both reflect the scope of each network as well as the extent to which each complex was found to be highly interactive.

#### Randomization of the BioPlex Network

Several analyses described below use randomized interaction networks to define null distributions against which score distributions derived from BioPlex 3.0 may be compared. One important consideration in randomizing the BioPlex network is that it contains nodes of two types – baits and preys – with slightly different properties. Most significantly, baits have greater numbers of interactions on average, compared to proteins detected only as preys. This feature must thus be maintained within the randomized network. During network randomization, total numbers of edges and nodes were maintained, and vertex degree was conserved for every protein. We further ensured randomized edges still connected baits (i.e. proteins targeted for AP-MS in BioPlex 3.0) with proteins in fact detected as preys. In other words, randomized edges were not allowed to connect pairs of proteins for which neither had been targeted for affinity purification. Furthermore, randomized edges were only allowed to connect pairs of baits if at least one of them was detected as a prey as well.

#### Subcellular Fractionation Correlation among BioPlex Interacting Proteins

Because proteins must at least partially co-localize to interact, we used subcellular fractionation profiles (‘Subcell Bar Codes’) for human proteins taken from a previously published study (Orre et al., 2019). Each protein in BioPlex 3.0 was matched with its subcellular fractionation profile. Then each edge in BioPlex 3.0 was scored by calculating the Pearson correlation coefficient for the subcellular fractionation profiles of its constituent proteins. If one or both proteins could not be matched with its subcellular fractionation profile, then that interaction was skipped. In parallel, edges of an equivalent randomized BioPlex 3.0 network were also scored. Because separate ‘Subcell Bar Codes’ were reported for four different cell lines – A431, H322, MCF7, and HCC827 – this entire procedure was repeated four times and Pearson correlation coefficients were averaged across cell lines for each edge in the real and randomized BioPlex networks. Distributions of real and random correlation coefficients were compared using a Cramér-von Mises test.

#### Size-exclusion Chromatography Elution Profile Correlation among Interacting Proteins

To evaluate the tendency of interacting proteins pairs in the BioPlex to co-fractionate during size-exclusion chromatography, we mapped previously published profiles (Heusel et al., 2019) obtained from fractionation of 293T lysate to proteins in the BioPlex 3.0 network. Each interaction was scored by calculating the Pearson correlation coefficient between co-elution profiles of its two constituent proteins. If one or both proteins did not have associated co-fractionation data, that interaction was skipped. In parallel, edges of a randomized BioPlex 3.0 network were also scored. Distributions of correlation coefficients were compared with a Cramér-von Mises test.

#### Graph Assortativity Enrichment among GO-SLIM Terms

To quantify the tendency for proteins of shared function to associate with each other in the BioPlex 3.0 network, graph assortativity was measured for GO-SLIM terms and compared to that observed across a population of randomized networks. GO categories were downloaded from www.geneontology.org in November 2018 and filtered to include only GO-SLIM terms. Each GO-SLIM term was mapped to BioPlex 3.0; if at least five proteins in the network matched the indicated category, it was scored to determine graph assortativity. This calculation was then repeated across 1,000 randomized networks and the resulting null distribution was used to determine a mean and standard deviation that could be used to convert the graph assortativity observed for that term in the real BioPlex 3.0 network into a Z-score. A Z-test was performed to calculate a p-value whose value was required to be smaller than 0.01 after Bonferroni correction for significance.

#### Analysis of BioPlex Reciprocal Interactions

Reciprocal interactions were tallied within BioPlex 3.0 and 1,000 randomized versions. Counts calculated from randomized networks were used to determine a mean and standard deviation for the null distribution and convert the reciprocal count observed in BioPlex 3.0 into a Z-score. A Z-test was subsequently performed to derive an associated p-value.

To determine the reciprocal rate observed in BioPlex 3.0, the total number of reciprocal interactions was compared against the number of edges that were eligible for reciprocal detection. To be eligible for reciprocal detection, an edge had to connect two proteins both of whom 1) were targeted for AP-MS as a bait; and 2) were detected as a prey in 293T cells.

#### Analysis of BioPlex Cliques

Cliques of three mutually interacting proteins (3-cliques) were tallied in BioPlex 3.0 using *Mathematica*. This process was repeated in 1,000 randomized networks as well. After calculating a mean and standard deviation for the resulting null distribution, a z-score was calculated for the observed 3-clique count in BioPlex 3.0 and a Z-test was performed to determine its associated p-value. The number of 3-cliques supporting each BioPlex 3.0 edge was determined by identifying first-degree neighbors shared by each edge’s two constituent proteins.

#### Analysis of 293T Replicates

To evaluate reproducibility of AP-MS in 293T cells, selected baits were targeted a second time for AP-MS. These replicate IP’s were performed under identical conditions and were scored against the same “stats” table used for generation of the 293T network with identical filtering to identify interacting proteins.

To determine the replication rate, each replicate IP was matched with the original pull-down of the same clone in the BioPlex 3.0 network (**Table S2E**). Those interactions resulting from pull-down of the target clone in the original BioPlex 3.0 network were extracted and each edge was labeled according to whether that interaction was also observed in the replicate IP. An interaction was considered replicated if it was 1) detected in the original BioPlex 3.0 IP among the top 2% of all bait-prey pairs in all 293T IP’s; and 2) was detected in the replicate IP with a score high enough to rank in the top 5% of all bait-prey pairs in all 293T IP’s. See **Table S2F** for a summary of interaction replication.

### Comparison of 293T and HCT116 Networks (Figures 2 & S2)

#### Quantitative Analysis of HCT116 and 293T Proteomes

Following normalization and scaling to report each protein’s TMT expression profile as a fraction of total observed signal, unpaired T-Tests were performed assuming unequal variance for each quantified protein. Differentially expressed proteins were defined based on an absolute log_2_ fold change greater than 0.5 coupled with a p-value smaller than 0.01 following multiple testing correction (Benjamini and Hochberg, 1995). Differentially expressed proteins were further grouped according to whether they were elevated in 293T or HCT116 cells and GO enrichment analysis was performed. GO enrichments were calculated using GO classifications downloaded from www.geneontology.org in November 2018 and used as background the set of all proteins quantified via TMT. Enrichment was calculated based on a one-sided hypergeometric test with subsequent multiple testing correction (Benjamini and Hochberg, 1995). See **Table S2A**.

#### Comparison of Bait Expression Levels

Relative bait expression levels were approximated in 293T and HCT116 cells by comparing numbers of spectral counts observed matching each bait in paired IP’s. For this comparison, 293T and HCT116 IP’s were aligned according to the precise clone used for AP-MS. A regression line was fit to the resulting scatter plot while fixing the intercept at zero. A sign test was used to determine whether the median PSM difference (PSM_293T_ – PSM_HCT116_) across baits is zero or not.

#### Analysis of AP-MS Replicates within and between Cell Lines

The 293T replication rate was determined as described above (*Analysis of 293T Replicates).* The same basic procedure was repeated to assess the replication rate between 293T and HCT116 cells. First, IP’s performed in both cell lines were aligned according to the specific clone used for AP-MS analysis. Then for each shared clone, interactions were extracted from the BioPlex 3.0 network that resulted from its pulldown. Each interaction was considered replicated if it was 1) detected in the original BioPlex 3.0 IP among the top 2% of all bait-prey pairs in all 293T IP’s; and 2) it was detected in the HCT116 IP with a score high enough to rank in the top 5% of all bait-prey pairs in all HCT116 IP’s.

In total, 72 replicate IP’s were also performed in HCT116 cells as part of normal AP-MS pipeline operation. Replication rates were calculated essentially as described above (*Analysis of 293T Replicates*). One complication of this is that when the final HCT116 network was assembled and more than one IP was available for a given clone, we systematically chose to include the IP with the larger number of interacting proteins in the final network. Thus, to avoid bias when calculating the HCT116 replication rates displayed in **Figure S2K**, we randomly selected which replicate would be used as the basis for comparison for each clone. See **Table S2E-J**.

#### Quantifying Effects of Bait Expression on Interaction Replication Rates

To assess the effects of bait expression on interaction replication across 293T and HCT116 cells, IP’s were aligned according to the specific clone used for bait expression and the replication of each interaction was assessed as described above (*Analysis of AP-MS Replicates within and between Cell Lines*). Baits were then binned by relative expression ratio along with their associated interactions. Interactions in each bin were subsequently tallied to determine the fraction of interactions confirmed across cell lines.

#### Quantifying Effects of Prey Expression on Interaction Replication Rates

To assess the effects of bait expression on interaction replication across 293T and HCT116 cells, IP’s were aligned according to the clone used for bait expression. Those successfully targeted in both 293T and HCT116 cells were identified and their resulting interactions extracted from the larger networks. Replication of each interaction was assessed as described above (*Analysis of AP-MS Replicates within and between Cell Lines*). Preys associated with these selected interactions were then matched with their relative expression levels in 293T versus HCT116 cells as measured via TMT (see *Quantitative Analysis of HCT116 and 293T Proteomes*). Interactions were binned according to each prey’s expression ratio and then the rate of replication was determined within each bin.

### Analysis of Shared and Cell-specific Interactions (Figure S3)

#### Quantification of Overlap among Edges Eligible for Detection in both Cell Lines

Because fewer AP-MS experiments have been completed in HCT116 cells, a significant fraction of interactions in the 293T network are unique to that network simply because the appropriate IP’s have not yet been performed in HCT116 cells. Such interactions that could only be detected in a single cell line were excluded by filtering interactions to include only those for which at least one constituent protein was targeted as a bait in both 293T and HCT116 cells. Note that the definition used here is less stringent than that described for **Figure S2**. In **Figure S2** we required matched clones targeting the same bait protein to pull down the same prey in each cell line. Here we do not necessarily require the same protein to have been targeted as a bait in both cell lines. For example, a hypothetical interaction between two proteins A and B would be considered eligible for detection in both cell lines when protein A was a bait in 293T cells only and protein B was a bait in HCT116 cells only; moreover, this interaction could be replicated in both cell lines if we detected the directed interaction A → B in 293T cells and the complementary directed interaction B → A in HCT116 cells. This relaxed definition allowed us to retain a larger fraction of the overall interaction space for the comparative analyses presented below.

#### Comparison of Network Properties for Shared and Cell-line-specific Edges

To determine whether interactions shared across cell lines were likely to reside in more central network locations, edge betweenness centrality was calculated for all edges in the combined 293T/HCT116 network using *Mathematica 12.0*. Edges were subsequently filtered to include edges detectable in both cell lines as described above and partitioned into 10 equal bins according to their betweenness centrality. Within each bin the fraction of edges observed in both cell lines was determined. Error bars represent bootstrapped 95% confidence intervals.

Similarly, local clustering coefficients were calculated for each interaction observed in the combined 293T/HCT116 network by extracting the subnetwork bounded by the first-degree neighbors of both constituent proteins and calculating its clustering coefficient. As before, edges detectable in both cell lines were partitioned into 10 equal bins according to these clustering coefficients and within each bin the fraction of edges observed in both cell lines was determined. Error bars represent bootstrapped 95% confidence intervals.

#### Essentiality and Interaction Overlap Between Cell Lines

To evaluate any relationship between protein essentiality and interaction overlap between cell lines, proteins in the combined 293T/HCT116 network were labeled as either ‘essential’ or ‘not essential’ according to specific definitions described below. Interactions eligible for detection in both cell lines were then extracted from the combined network and binned according to whether 0, 1, or 2 of their constituent proteins met the indicated definition for essentiality. Finally, edges in each bin were labeled as ‘shared’ or ‘cell-line specific’ and the fraction of edges shared was calculated. Bootstrapped 95% confidence intervals were calculated as well.

This analysis was repeated three times, each time defining ‘essential’ genes according to separate datasets. First, we followed the criteria described by Wang et al. and deemed proteins with gene-based CRISPR scores < 0.1 and corrected p-value < 0.05 in KBM7 cells as ‘essential’ (Wang et al., 2015). Second, we counted as ‘essential’ all genes identified by Blomen et al. as fitness-associated in either KBM7 or HAP1 cells (Blomen et al., 2015). Third, we used ‘common essential’ genes as derived previously (Dempster et al., 2019) from Achilles data (Meyers et al., 2017; Tsherniak et al., 2017).

#### Interaction Overlap and Protein Expression Variability

In **Figure S2**, we observed that differential protein expression contributes to cell-line specificity among protein interactions. Expanding on this point, we wanted to see whether proteins involved in cell-line-specific interactions are variably expressed more generally across other biological contexts. For this purpose, we obtained datasets reporting protein expression across 378 human cancer cell lines (Nusinow et al. *in press*) and gene and protein expression across 29 human tissues (Wang et al., 2019). Proteins in the cancer cell line dataset were ranked according to expression variability as measured by each protein’s standard deviation across cell lines; similarly, proteins and genes in the human tissue datasets were ranked according to relative standard deviations observed across tissues. In each case, proteins or genes were partitioned into 10 equal bins. Interactions eligible for detection in both cell lines were then extracted from the combined 293T/HCT116 network and binned according to the ranked expression variability of each constituent protein. Finally, within each bin the fraction of edges observed in both cell lines was determined.

#### Interaction Overlap and PubMed Citations

All proteins in the combined 293T/HCT116 network were linked with their associated PubMed citation counts as downloaded via FTP from NCBI in December 2018. Proteins in the network then partitioned into 10 equal bins ranked by citation count. Interactions eligible for detection in both 293T and HCT116 cells were then partitioned according to the citation bins of their constituent proteins. Finally, the overlap rate was calculated for all interactions in each partition.

### Evolutionary Analysis of BioPlex Edges (Figure S4)

To assess the evolutionary context of shared and cell-line-specific interactions, proteins within the combined 293T/HCT116 network were matched with protein evolutionary age data as published previously (Liebeskind et al., 2016). Following the authors’ guidelines, we only accepted protein age estimates with the following parameters: entropy < 1.0, hgt_flag = false, numDBsContributing > 3, and Bimodality < 5. Proteins without evolutionary ages from this study meeting these criteria were excluded from this analysis. Each evolutionary split was assigned with an approximate date using TimeTree.org. The category Eukaryota + Bacteria was assigned the midpoint between dates associated with the emergence of Eukaryotes and Opisthokonts.

In some contexts (e.g. **Figure S4D**), interactions were binned according to the evolutionary ages assigned to both of their constituent proteins; however, in other situations (e.g. **Figure S4A/C**) it was necessary to assign a single evolutionary age to each edge. In these cases, edges were assigned the age of their younger constituent protein. Edges for which one or more proteins could not be assigned ages were excluded from this analysis.

In these analyses, edges were deemed eligible for detection in both cell lines according to the same criteria described above for **Figure S3** (*Quantification of Overlap among Edges Eligible for Detection in both Cell Lines*).

### Analysis of Functionally Defined Subnetworks (Figure 3)

To determine the extent to which functionally defined subnetworks of the human interactome were conserved across 293T and HCT116 cell lines, we extracted subnetworks corresponding to a wide range of functionally defined categories. These include CORUM (Giurgiu et al., 2018) complexes (coreComplexes.txt; downloaded from https://mips.helmholtz-muenchen.de/corum/ in September 2018); Reactome (Fabregat et al., 2017) pathways (downloaded via www.uniprot.org in January 2019); GO Biological Process, Cellular Component, and Molecular Function (Ashburner et al., 2000) categories (www.geneontology.org; November 2018); and DisGeNET (Piñero et al., 2017) disease associations (www.disgenet.org; May 2019).

When calculating overlap of interactions within subnetworks for this analysis, interactions were only counted if at least one of their constituent proteins was targeted for AP-MS analysis in both cell types. This corrects for the fact that some interactions are not detectable in one cell line or the other simply based on the baits targeted in each. However, when subgraphs were plotted in the figures, all matching proteins and interactions were displayed including edges that could only be detected in a single cell line. Results are summarized in **Table S3**.

### Discovery of Shared and Cell-line-specific Network Communities (Figure 4)

The combined 293T/HCT116 interaction network was partitioned into communities as described previously (Huttlin et al., 2017). Briefly, MCL clustering (Enright et al., 2002) was used to subdivide the network into communities using an inflation parameter of 2.25 and setting the ‘force-connected’ option to yes. Communities with fewer than 3 members were discarded, leaving 1,423 communities overall (**Table S4A-B**).

After subdividing the combined 293T/HCT116 interaction network into communities, we next sought pairs of communities whose members were found to interact with unusually high frequency, as described previously (Huttlin et al., 2017). First, the full set of interactions was trimmed to include only those interactions connecting one community to another. For each pair of communities connected by one or more edges, the numbers of edges emanating from each were determined, as were the number of edges connecting the two. Fisher’s Exact Test was then used to assess whether edges connecting the two were enriched beyond random chance, followed by multiple testing correction (Benjamini and Hochberg, 1995). A total of 1,736 statistical associations were detected between communities at a 1% FDR (**Table S4C**).

Having identified communities and community associations in the combined 293T/HCT116 network, we next wanted to quantify the level of overlap observed for each among cell lines. To do this, we first filtered our list of communities down to 761 that contained at least one protein which had served as an AP-MS bait in both cell lines. Within each of these we assessed the overlap as discussed below (**Table S4D**). Similarly, we filtered 1,736 interactions down to a subset of 988 community-community associations ensuring that at least three interactions bridging these communities were detectable in both cell lines (**Table S4E**).

When calculating overlap within communities or between communities, interactions were only counted if at least one of their constituent proteins was targeted for AP-MS in both cell types. This accounts for interactions detectable in only a single cell line simply due to the specific AP-MS experiments completed in each. However, **Figure 4E-H** include all matching proteins and interactions, including those only eligible for detection in a single cell line.

### Analysis of Consistency of Function among Interactors (Figure S6)

Enrichment analysis was performed to assess consistency of function among each protein’s first-degree neighbors. For each protein in the combined 293T/HCT116 network, we extracted all first-degree neighbors. Functional categories including GO-SLIM Biological Process, Molecular Function, and Cellular Component categories (www.geneontology.org; November 2018), PFAM domains and Reactome pathways (downloaded via www.uniprot.org; January 2019) were superimposed. Enrichment of each term within a protein’s first-degree neighbors was calculated using a one-sided hypergeometric test with multiple testing correction (Benjamini and Hochberg, 1995). Those significant at an FDR less than 1% were retained.

For every functional category deemed significant after enrichment analysis, those first-degree neighbors matching the category were assessed to determine what fraction were significant to 293T cells, significant to HCT116 cells, or common to both. At least five neighboring proteins were required to match a given functional category to be included in **Figure S6A**. Results are provided in **Table S5C**.

### Domain-domain Associations across Cell Lines (Figure 5)

First, the complete set of interactions in the combined 293T/HCT116 network was extracted and mined for associations among protein domains as described previously (Huttlin et al., 2015). To start, all proteins in the network were linked to PFAM domains (El-Gebali et al., 2018) as recorded in Uniprot (downloaded in January 2019). Domain pairs connected by two or more interactions were assessed for significance using Fisher’s Exact Test after accounting for 1) the number of interactions connecting both domains; 2) the numbers of interactions involving either domain individually; and 3) the number of interactions not involving either domain. After multiple correction (Benjamini and Hochberg, 1995), those domain-domain associations significant at a 1% FDR were identified (**Table S5A**).

Second, each statistically significant PFAM domain association was assessed for overlap between cell lines (**Table S5B**). For each pair of associated domains, all corresponding interactions were identified. This list of interactions was then filtered to include only interactions detectable in both cell lines, in the sense that at least one member of each interacting protein pair must be a bait in both cell types. If at least five edges remained, these were then tallied in both cell lines to determine the proportions specific to a single cell type or shared across both. These 3,249 domain-domain associations were also assembled into a network whose nodes corresponded to PFAM domains and whose edges linked statistically associated PFAM domain pairs.

### Assignment of Accessory Proteins to Known Protein Complexes (Figure 6)

New proteins were associated with known complexes based on statistical enrichment. We first describe the general approach as applied to a single network before explaining how it was extended to incorporate two independent interaction networks.

To start, a set of proteins constituting a known protein complex was mapped onto an interaction network. First-degree neighbors of these known complex members were identified as candidate complex members whose association was tested statistically. This statistical association was tested via Fisher’s Exact Test, taking into account 1) the number of interactions linking the candidate complex member to known complex members; 2) the number of interactions the candidate complex member makes to other unrelated proteins; 3) the number of interactions that connect known complex members to other proteins; and 4) the number of interactions that involve neither known complex members nor the putative complex member being scored. Note that this accounting overlooks interactions that connect known complex members to each other.

To leverage the availability of two independent interaction networks, the procedure described above was modified slightly. First, each known complex was mapped onto both 293T and HCT116 networks and any protein found adjacent to one or more known complex members in either network was considered a putative complex member. Each was then scored independently using either the 293T network or the HCT116 network as described above to produce two p-values. These p-values were then combined using Fisher’s Method to derive a single p-value reflecting the combined statistical strength of the evidence linking each putative complex member with the known complex.

In either case, whether a single network or multiple networks were used for prediction, the final set of p-values was adjusted for multiple testing correction (Benjamini and Hochberg, 1995) and an adjusted p-value smaller than 0.01 was required for significance.

In principle, known complexes could come from a range of sources. We targeted non-redundant CORUM (Giurgiu et al., 2018) complexes (coreComplexes.txt; downloaded in September 2018). Since leave-one-out cross-validation was to be applied, a couple of additional filters were applied. At least four members were required overall, and at least four members had to be present in the combined BioPlex network. In addition, at least three of these proteins had to be targeted as baits. Finally, when multiple entries matched the same complex (as given by the CORUM complex name), we kept the largest version only. This reduced set of complexes was used both for leave-one-out cross-validation and for identification of additional complex members (**Table S6A-B**).

In addition to scoring CORUM complexes as described above, this method can also be used to score user-defined complexes. The tubulin polyglutamylase complex shown in **Figure 6H** is a user-specified complex that illustrates this point. Results are provided in **Table S6C**.

### Identification of Functional Associations using Achilles Data (Figure 7)

To map functional relationships onto the combined BioPlex network, gene fitness profiles defined via CRISPR knockout or RNAi knockdown across hundreds of cell lines were superimposed onto the network. Fitness profiles were originally collected through Project Achilles (Meyers et al., 2017; Tsherniak et al., 2017) and were downloaded through their public data portal in May 2019. For every pair of interacting proteins in the combined BioPlex network, fitness profiles were sought. If fitness profiles were available for both interacting proteins, their functional similarity was assessed using Spearman’s Correlation; statistical significance was assessed using the CorrelationTest function in *Mathematica 12.0* with subsequent multiple testing correction (Benjamini and Hochberg, 1995). Those BioPlex interactions found to match correlated fitness profiles at a 5% FDR were split into groups according to whether the observed correlation was positive or negative. Edges were also labeled to indicate whether they were observed in 293T cells, HCT116 cells, or both. Subnetworks corresponding to BioPlex interactions with either positive or negative fitness correlations were extracted and plotted using Gravity Embedding.

### DATA AND CODE AVAILABILITY

This project has generated many types of data and code that are available for distribution via numerous venues. Items not listed here will be provided from the Lead Contact upon reasonable request.

First, both 293T and HCT116 networks are described in full in the supplementary tables included with this manuscript, as are results of all analyses described herein. These include functional and disease associations, predicted localizations, communities, PFAM domains, replicate analyses, etc. See supplementary table legends for details.

Second, both filtered and unfiltered lists of interactions in HCT116 and 293T networks are available for download on the BioPlex website at bioplex.hms.harvard.edu/downloadInteractions.php. These interactions may be accessed either as flat files or through a custom API.

Third, both 293T and HCT116 networks, as well as the combined BioPlex network, will be deposited into NDEx (https://home.ndexbio.org/index/), the repository for biological network data (Pratt et al., 2015).

RAW files corresponding to both 293T and HCT116 AP-MS experiments may be accessed in multiple ways. First, if RAW files corresponding to a small number of specific baits are desired, these may be downloaded from the BioPlex website via a search interface at https://bioplex.hms.harvard.edu/downloadData.php. If larger numbers of RAW files are desired, they will be accessible via the MassIVE repository (ftp://massive.ucsd.edu). Finally, all ∼30,000 RAW files corresponding to 293T and HCT116 AP-MS networks are available upon request via GLOBUS infrastructure.

CompPASS software is available as an R package (https://github.com/dnusinow/cRomppass); we also make it available for small to medium-scale AP-MS experiments online via the BioPlex website: https://bioplex.hms.harvard.edu/comppass/.

RAW files corresponding to the RTS-MS3-TMT comparison of 293T and HCT116 proteomes have been deposited in MassIVE (ftp://massive.ucsd.edu).

### ADDITIONAL RESOURCES

In addition to the numerous resources described previously, we have also developed a full-featured web-based viewer, BioPlexExplorer, as a companion to this paper. For best results, we recommend using this tool in Chrome, though the full range of browsers are supported, including mobile browsers. This tool, available at bioplex.hms.harvard.edu/bioPlexExplorer, provides numerous features.

1. An integrated search engine for locating and viewing shared and cell-specific networks centered on one or more user-selected target proteins. All interactions are displayed as a fully interactive network diagram. Additional data, including evolutionary age, protein expression levels, and GO category membership may be superimposed onto the networks. These networks may be downloaded as images (png) or as Cytoscape networks (json). All interactions are also displayed in sortable tabular format and may be exported in Excel, CSV, or PDF formats. The resulting network viewer also performs GO enrichment analysis on-the-fly. Finally, these network views may be shared with colleagues and collaborators via custom URLs.
2. A custom ternary plot viewer that enables the user to view interactive versions of the ternary diagrams shown in **Figure 3A/E** as well as **Figure 4C**. Through a drop-down menu the user may select specific annotation sets of interest, including GO Cellular Component, Molecular Function, and Biological Process, Reactome Pathways, CORUM Complexes, DisGeNET diseases, and BioPlex communities. By zooming in on these ternary plots and clicking on specific points, the user may see cell-line-specific interaction data for thousands of complexes, communities, and functional categories. Links also enable the user to easily open a network view corresponding to each category of interest as well.
3. A fully interactive version of **Figure 4B**, which depicts the full set of communities observed in the combined BioPlex network. Statistically significant associations among BioPlex communities are displayed as well. This interface is fully searchable, enabling a user to quickly and easily locate proteins of interest; moreover, upon selecting a node, a link is available to easily view the underlying interaction data in both 293T and HCT116 networks.
4. A fully interactive version of the PFAM Domain Association Network depicted in **Figure 5A**. Nodes correspond to specific PFAM domains while edges correspond to pairs of domains that associate statistically. The color of each edge indicates the extent that each domain association is reflected in 293T and HCT116 cells individually or in common. Clicking on individual nodes reveals lists of proteins that possess each domain; in addition, a link is provided to view the interaction subnetwork defined by proteins with the indicated domain.
5. Fully interactive versions of **Figure 7B/C**, depicting BioPlex subnetworks that exhibit either positive or negative fitness correlations. Again, these networks are fully searchable and interactive.

## SUPPLEMENTARY TABLES

**Figure S1:**
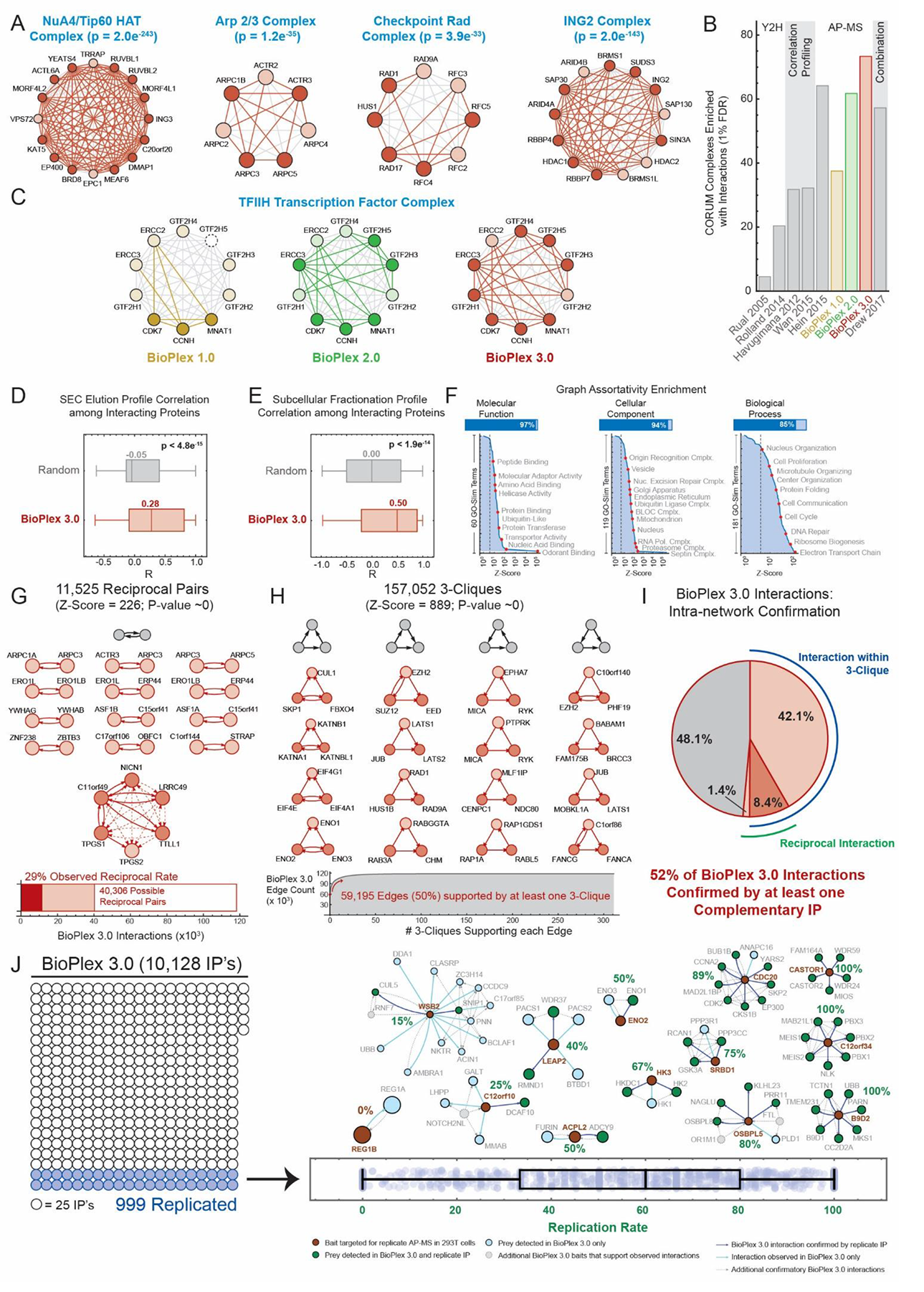
Validation of the BioPlex 3.0 Network, Related to Figure 1. (A) – (C) BioPlex 3.0 validation against CORUM complexes. To quantify whether each CORUM complex is reflected in each interaction network, the subnetwork defined by its constituent proteins was extracted from each network and the enrichment of interactions calculated relative to global network density. Focusing on enrichment, rather than interaction counts, is useful when comparing networks of variable size: smaller networks with fewer edges have lower interaction probabilities, so fewer edges are required to attain significance; in contrast, a larger network with more interactions has a higher probability of interaction and the number of interactions required for significance increases accordingly. (A) CORUM complex coverage within the BioPlex 3.0 network. Subnetworks corresponding to four CORUM complexes are displayed. Deep red nodes were targeted as baits while light red nodes were detected as preys only. Interactions observed via AP-MS are highlighted in red. (B) We show the fraction of CORUM complexes that were found to be significantly enriched for protein interactions (1% FDR) in nine different interaction datasets. Datasets are grouped according to the experimental methods used to create each. Note that some CORUM complexes may not be detectable in one or more AP-MS datasets because no complex members were targeted for AP-MS. No correction was applied because it would complicate comparisons across datasets, especially using different experimental approaches. (C) TFIIH Transcription Factor Complex coverage across BioPlex networks. Dark-colored nodes were targeted in baits in each network while light-colored nodes were present only as preys. Nodes with dashed outlines were not detected in the network. Observed interactions are colored. (D) To assess the tendency for interacting proteins to co-fractionate, size exclusion chromatography (SEC) elution profiles (Heusel et al. 2019) were correlated for all interacting proteins in BioPlex. Distributions of correlation coefficients are displayed for both real and randomized networks, as above. P-value: Cramer-von Mises test. (E) To assess the tendency for interacting proteins to co-localize, subcellular fractionation profiles (Orre et al. 2019) were correlated for all interacting protein pairs in BioPlex. In tandem, we repeated the analysis for a randomized version of the BioPlex network. Distributions of correlation coefficients are displayed for both real and randomized networks. P-value: Cramer-von Mises test. (F) The tendency for functionally related proteins to co-associate was measured by calculating the graph assortativity for each GO term. Calculations were then repeated across 1,000 randomized networks to estimate null distributions and Z-scores and p-values calculated via a Z-test with multiple testing correction. Z-scores are plotted for each GO-Slim term. Selected terms are labeled. Bar graphs indicate the fraction of GO-Slim terms in each category that significantly associate at a 1% FDR. (G) – (I) Intra-network validation of BioPlex 3.0 interactions. (G) To be validated reciprocally, both interacting proteins must have been targeted as baits and must be detectable as preys in 293T cells. A bar chart depicts the fractions of BioPlex edges either observed or eligible for reciprocal detection. In addition, several reciprocal pairs are displayed. The Z-score and p-value reflect enrichment of reciprocal pairs compared to 1,000 randomized networks (see Methods). (H) The lower cumulative histogram indicates the fraction of BioPlex edges that participate in one or more 3-cliques. Several 3-cliques are shown matching the 4 possible 3-clique topologies. The Z-score and p-value reflect enrichment of 3-cliques compared to 100 randomized networks (see Methods). (I) Pie chart depicting BioPlex edges confirmed by either reciprocal interactions or 3-cliques. (J) To assess reproducibility, we repeated AP-MS analysis of 999 bait proteins in 293T cells. The schematic on the left highlights the fraction of BioPlex 3.0 baits repeated for validation; the box-whisker plot at the bottom displays the fraction of interactions that were confirmed upon replication for each bait. A subset of these interaction networks are displayed above the box-whisker plot in positions that approximately correspond to their observed replication rate (written in green). Replicated baits are red; replicated preys are green while preys detected in the initial IP only are light blue. Replicated edges are green while edges only detected in the initial IP are light blue. Gray nodes and dotted edges depict additional confirmatory interactions in the BioPlex 3.0 network.

**Figure S2:**
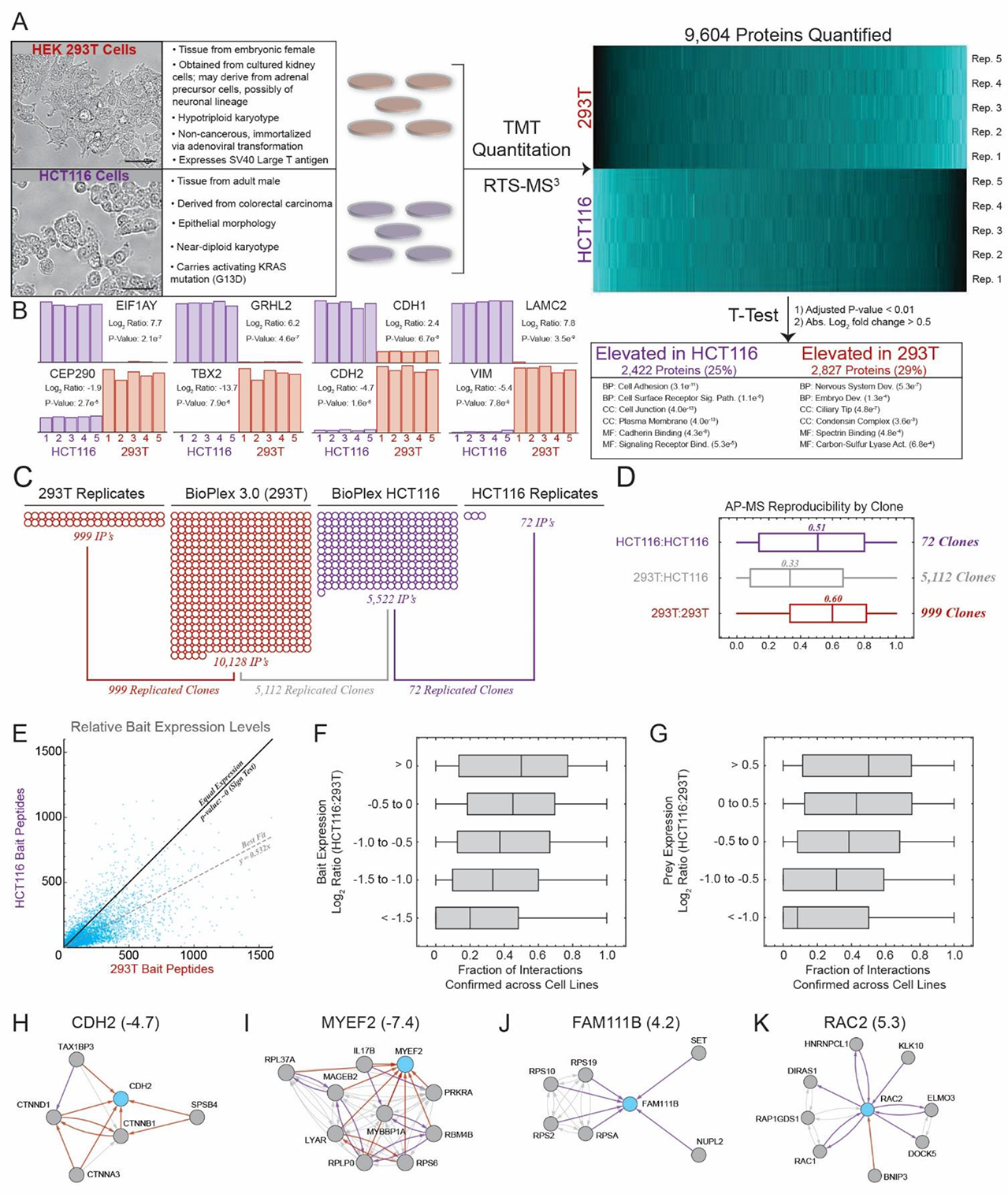
293T and HCT116 Cells Afford Complementary Perspectives on the Human Proteome and Interactome. Related to Figure 2. (A) 293T and HCT116 cell lines arise from very different sources and exhibit distinct phenotypes as summarized in the table. To explore how these differences manifest within their proteomes, we compared biological quintuplicates of each cell line using TMT isobaric labeling coupled with Real-Time Search MS^3^ quantitation (Erickson et al. 2019). Protein expression profiles are summarized in the heatmap at the right; GO enrichment analysis was applied to proteins elevated in each cell line. Enriched GO terms are summarized in the table below. (B) TMT profiles observed for selected cell-line-specific proteins. Reporter ion intensities have been normalized so that each TMT profile sums to 1.0 across all channels. P-values were calculated with T-Tests and corrected for multiple testing. (C) Schematic depicting AP-MS analyses and replicates performed in 293T and HCT116 cells. Note that the numbers of clones targeted in both 293T and HCT116 cells differ slightly from the numbers of shared bait proteins listed in Figure 2C. The ORFeome includes multiple clones that match some human genes. Whenever possible, the same clones were used for each bait in both cell lines; however, in a small number of cases AP-MS analysis of the preferred clone failed in HCT116 cells while another clone matching the same gene was completed successfully and incorporated into the final network. When viewed at the gene/protein level as in Figure 2C, these would be counted as shared between cell lines. However, when compared as the clone level as here, these would not be counted as shared. (D) Box-whisker plot depicting bait-specific replication rates observed within and across cell lines (E) Scatter plot comparing abundances observed in 293T and HCT116 cells for baits targeted in both cell lines. A solid black line denotes equal expression; data points show a significant tendency to fall below this line (p ∼ 0; Sign Test). A dashed gray line indicates the best fit calculated while fixing the intercept at zero. (F) Interaction confirmation versus bait expression. Baits targeted in both cell lines were assigned to 5 bins according to relative bait expression as reflected by numbers of bait peptides in each IP. Box-whisker plots depict fractions of interactions confirmed for baits in each bin. (G) Interaction confirmation versus prey expression. Baits targeted in both cell lines were extracted from the larger interaction networks along with their associated prey proteins. These interactions were then divided into bins based on prey expression as measured by TMT. Box-whisker plots depict fractions of interactions confirmed for preys in each bin. (H) – (K) Interaction networks of selected cell-line-specific proteins. Target proteins are named at the top of each network and are highlighted in blue; neighboring proteins are gray. Edges observed in both cell lines are gray while 293T-specific edges are red and HCT116 edges are purple. Directed edges point from baits toward preys. Numbers in parentheses after each protein are the log2 ratios of expression in HCT116 cells relative to 293T cells.

**Figure S3:**
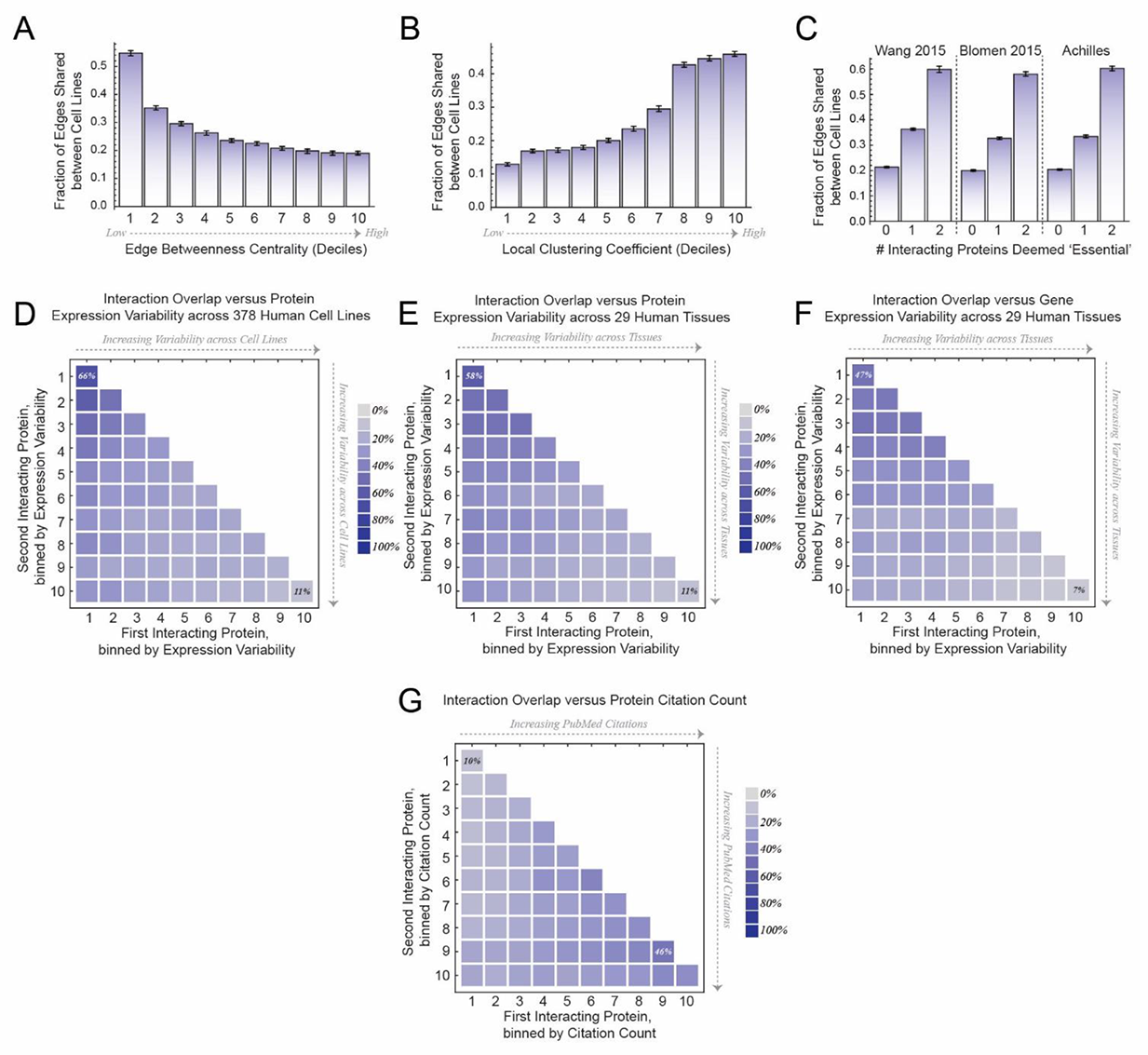
Distinct Properties of Shared and Cell-line-specific Interactions. Related to Figure 2. (A) – (B) The combined 293T/HCT116 network was filtered to include only edges eligible for detection in both cell lines (i.e. at least one protein in each interacting pair must be a bait in each cell line). These edges were then divided into deciles based on edge betweenness centrality (A) or local clustering coefficient (B) and the fraction of shared edges in each decile was calculated. Error bars: bootstrapped 95% confidence intervals. (C) Edges detectable in both cell lines were grouped according to whether 0, 1, or 2 of their constituent proteins were classified as ‘essential’ or associated with increased cellular fitness in datasets from Wang et al. 2015, Blomen et al. 2015, or the Achilles project (Dempster et al. 2019). The edge overlap between cell lines was calculated for each class. Error bars: bootstrapped 95% confidence intervals. (D) – (F) Edges detectable in both cell lines were binned according to the relative variability of expression observed for their constituent proteins across 378 human cancer cell lines (Nusinow et al. 2019) (D); expression patterns of proteins (E) and genes (F) across 29 human tissues were considered as well (Wang et al. 2019). The fraction of edges shared between cell lines was calculated for each bin. Bins with maximum and minimum overlap are labeled. (G) Edges eligible for detection in both cell lines were binned according to the numbers of PubMed citations for their constituent proteins and the fraction of edges shared between cell lines was calculated for each bin. Bins with maximum and minimum overlap are labeled.

**Figure S4:**
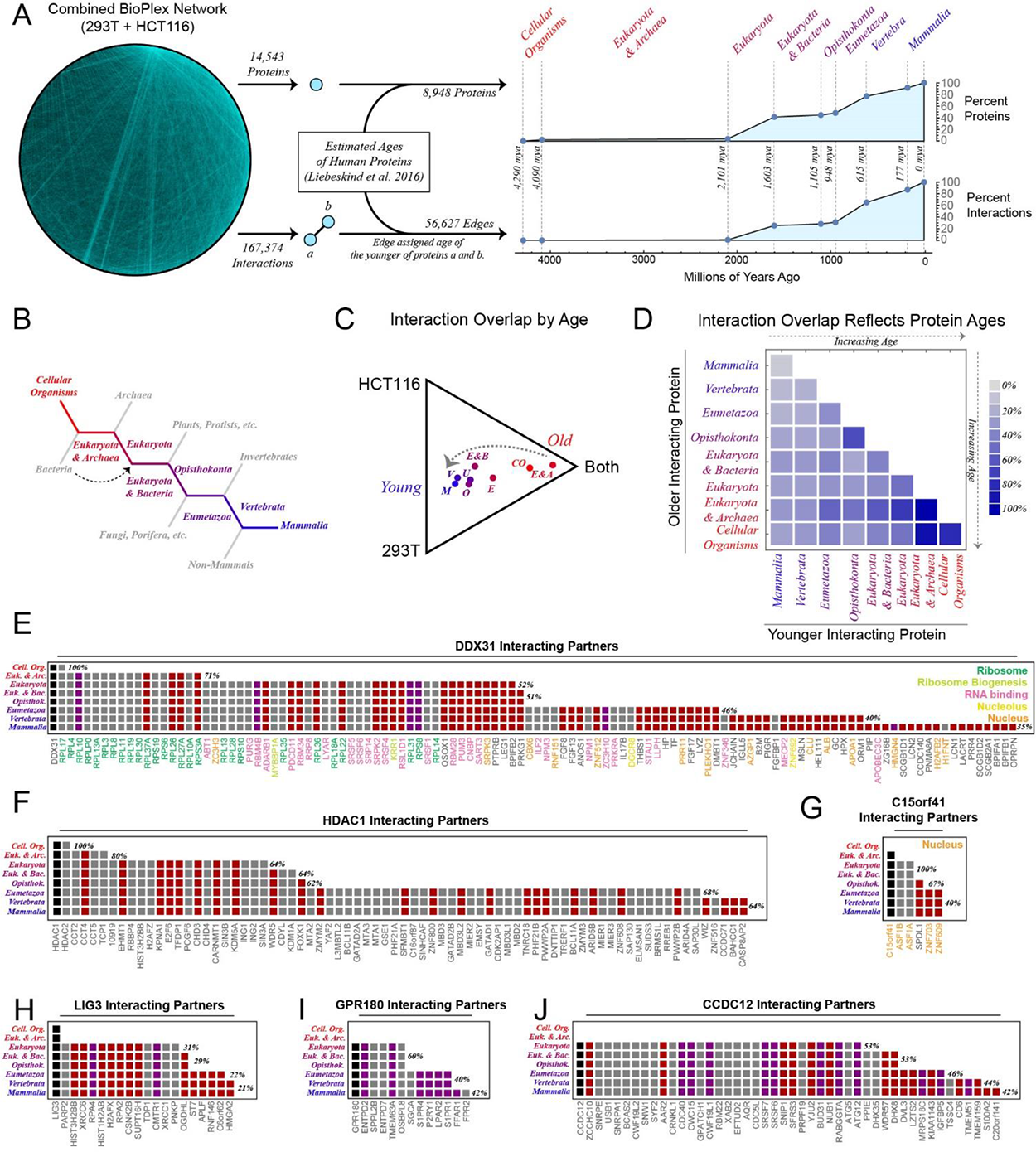
Interactions Shared across Cell Lines Correlate with Evolutionary Age. Related to Figure 2. (A) Consensus evolutionary ages from Liebeskind et al. 2016 were mapped onto the combined BioPlex network to assign ages to individual proteins; interactions were then assigned the age of their youngest constituent protein. Line plots show the fractions of BioPlex proteins and interactions that can be traced to each of 8 ages. Approximate dates for each split were taken from TimeTree.org. Only proteins whose evolutionary ages could be estimated with sufficient confidence were included in this plot (see Methods); likewise, interactions are only included if both constituent proteins could be assigned evolutionary ages. Interactions were further filtered to include only those involving at least one protein targeted for AP-MS in each cell line. Evolutionary ages assigned to individual proteins and interactions may be viewed at **bioplex.hms.harvard.edu/bioPlexExplorer**. (B) Simplified diagram showing phylogenetic relationships among assigned evolutionary ages. (C) Interactions were binned according to estimated evolutionary age of their youngest constituent and filtered to include only those interactions that were detectable in both cell lines (i.e. involved at least one protein targeted as a bait in each cell line). The proportions of edges shared and unique to each cell line were then calculated within each bin. These proportions for each evolutionary age are plotted as points on a ternary diagram. The arrow highlights the progression toward higher cell-type specificity among interactions with more recent evolutionary origins. (D) Interactions were again binned, this time accounting for the evolutionary ages of both older and younger members. Within each bin the fraction of edges shared between cell lines was calculated. This fraction is plotted on the heat map as a gradient from light to dark blue. Interaction overlap rates ranged from 6 – 86%. (E) – (J) Heat maps depicting the evolutionary ages of interacting partners for selected BioPlex proteins. Target proteins are labeled at the top of each plot and are depicted in the leftmost column of each heat map. Proteins identified as interactors are then displayed in decreasing order of age. Each column in the heat map is colored to indicate whether the associated interacting partner was observed in 293T cells (red), HCT116 cells (purple), or both (gray). Percentages to the right of each row indicate the overlap among interactions dating to the indicated age or before. In some cases, bait names are colored to encode membership in specific functional categories or subcellular compartments.

**Figure S5:**
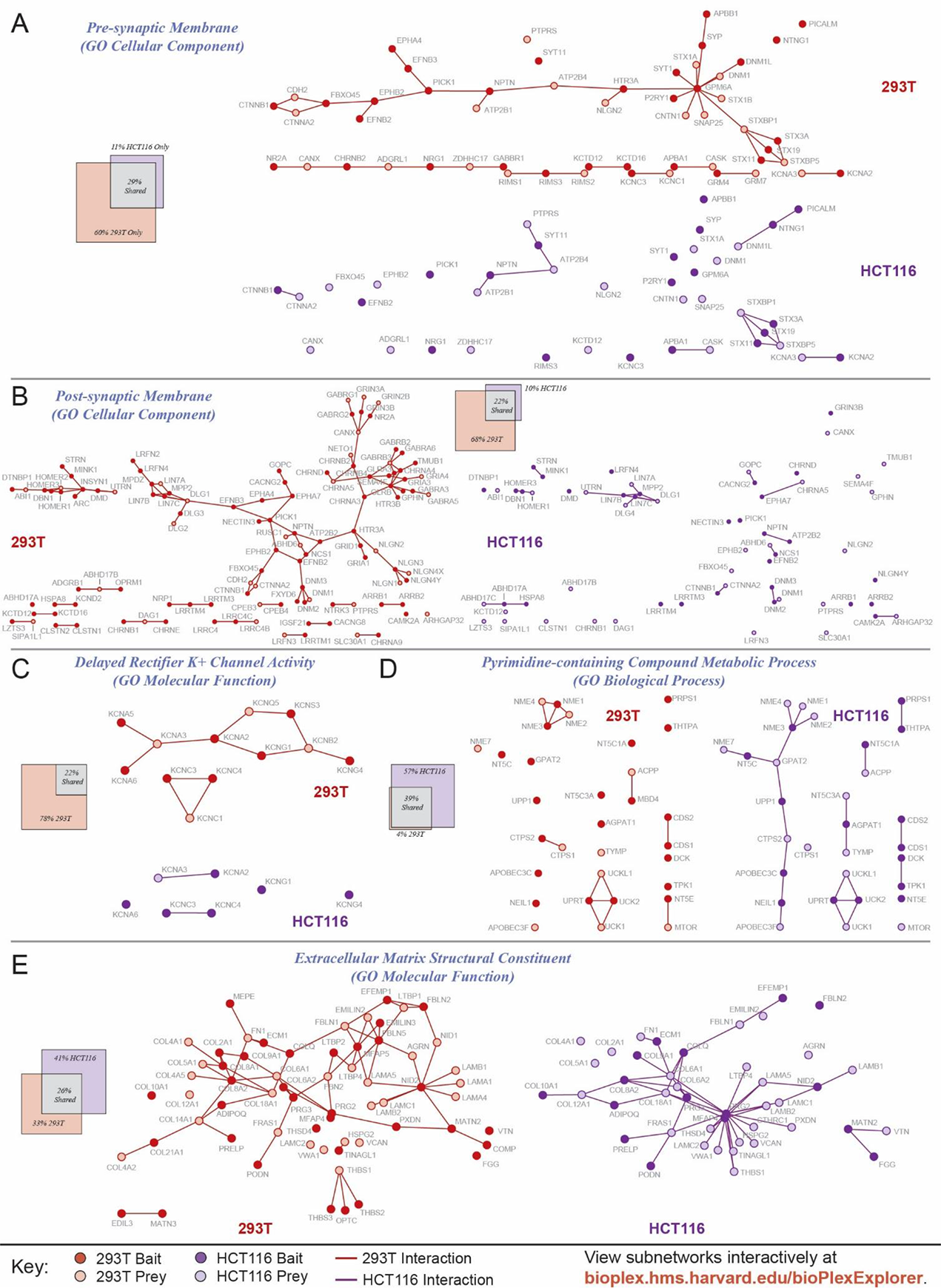
Functionally Defined Subnetworks are Shared Variably across Cell Lines: Additional Examples. Related to Figure 3. (A) – (E) Subnetworks corresponding to five GO categories are highlighted. 293T subnetworks are red while HCT116 subnetworks are purple. Baits targeted for AP-MS in each cell line are shaded. Layouts of the cell-line-specific networks are identical, except proteins missing from either network are absent. Each panel is accompanied by a Venn Diagram depicting the edge proportions shared and specific to either cell line. Additional subnetworks corresponding to GO categories and Reactome pathways may be viewed at **bioplex.hms.harvard.edu/bioPlexExplorer**.

**Figure S6:**
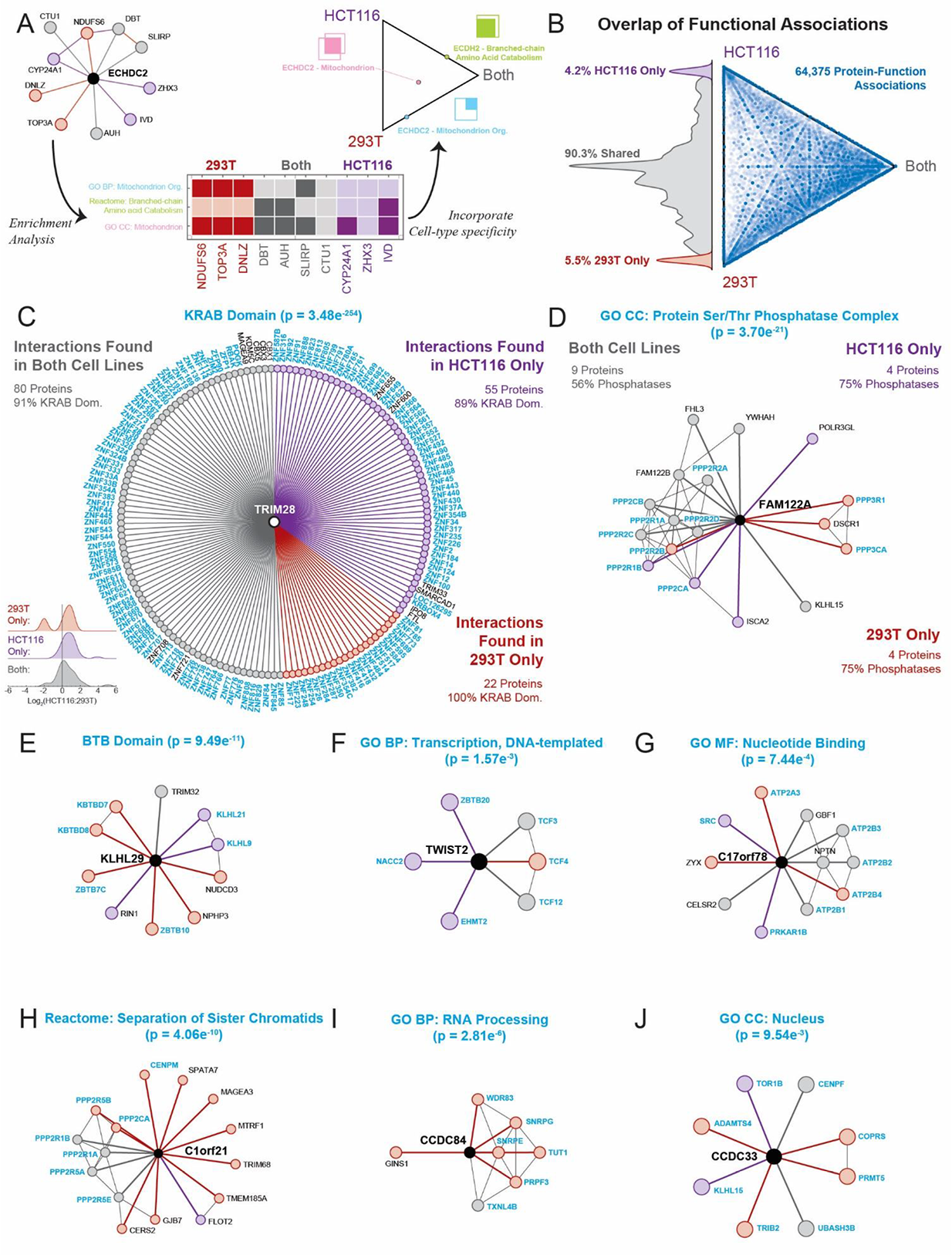
293T and HCT116 Interaction Profiles: Shared Function Despite Cell-Specific Interactions. (A) To evaluate consistency of biological function, individual proteins were selected from the combined 293T/HCT116 network along with their first-degree neighbors. Only proteins targeted as baits in both cell lines and found to bind at least one protein in each network were considered. For each protein, GO categories, Reactome pathways, and PFAM domains enriched among its primary neighbors were identified. As an example, the neighborhood surrounding ECHDC2 is displayed at the upper left; the target protein is black while shared nodes and edges are gray; those specific to 293T cells and HCT116 cells are red and purple, respectively. Among first-degree neighbors of ECHDC2 three terms were enriched (adjusted p < 0.01): Mitochondrion, Mitochondrial Organization, and Branched Chain Amino Acid Catabolism. These terms map to the primary neighbors as depicted in the central heat map: dark shading indicates that a prey matches the indicated classification. Within each category (i.e. row of the table) the relative proportions of edges shared or specific to a particular cell line can be tallied; each protein-function association can then be plotted within a ternary diagram as shown. (B) Ternary diagram depicting the extent to which each association between bait and biological function is observed in both cell lines. This ternary diagram is accompanied by a histogram that shows the cell line specificity of these terms after effectively projecting the individual points onto the vertical edge of the ternary diagram. This graphically depicts the extent to which these bait – function associations are shared or cell-line-specific. To be included in this plot, at least 5 neighboring proteins had to match the indicated functional category and category had to be enriched at a 1% FDR. (C) TRIM28 – KRAB domain functional association. Shown here are the interacting partners observed for the protein TRIM28 in both 293T and HCT116 cell lines. Nodes and edges are colored according to whether the interaction was observed in 293T cells (red), HCT116 cells (purple) or both (gray). Interactor names have been colored to indicate whether each contains a KRAB domain. The associated p-value reflects the enrichment of KRAB domains among first-degree neighbors of TRIM28. Inset histograms reflect the relative abundances of these TRIM28 interactors in 293T and HCT116 cells as measured by TMT (see **Figure S2**). (D) – (J) Additional protein – function associations. Each panel depicts a selected target protein (in black) along with its first-degree neighbors, colored to indicate whether the interaction was observed in 293T cells (red), HCT116 cells (purple) or both (gray). Labels of interacting partners that match the indicated functional category or bear the indicated domain are highlighted in blue. Interactions that do not involve the target proteins are rendered as thin, black lines. P-values reflecting the strength of the protein – function association are displayed along each title.

**Figure S7:**
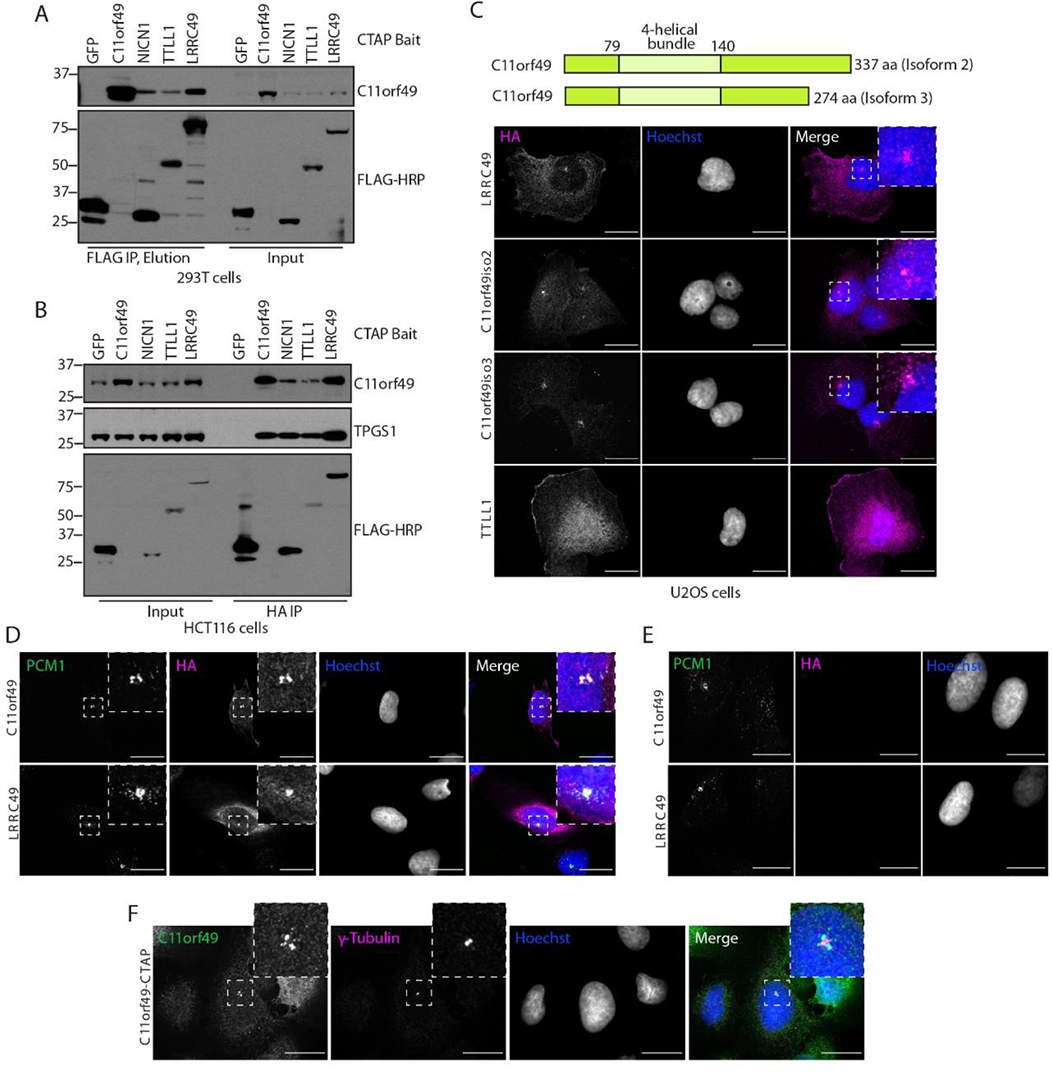
C11orf49 Associates with the Tubulin Polyglutamylase Complex. Related to Figure 6. (A) – (B) Endogenous C11orf49 binds to members of the TTLL1 polyglutamylase complex.C-terminally FLAG-HA tagged TTLL1 complex proteins or GFP were expressed in 293T or HCT116 cells followed by FLAG or HA purification and western blotting with endogenous C11orf49 antibodies. (C) Localization of TTLL1 complex members and C11orf49 long and short isoforms. The longest C11orf49 isoform (isoform 2) along with a truncated isoform (isoform 3) were expressed in U2OS cells. Cells plated on high precision coverslips were fixed and immunostained with anti-HA antibodies to visualize ectopic protein localization via confocal microscopy. Max intensity projections of 2-µm z-stacks are shown; scale bar represents 20 µm. (D) C11orf49 exhibits pericentriolar localization. U2OS cells stably expressing C11orf49 or TTLL1 complex member LRRC49 were fixed, stained with anti-HA and anti-PCM1 (pericentriolar marker) antibodies and visualized by confocal microscopy. Single planes of a z-stack are shown; scale bar represents 20 µm. (E) Staining with PCM1 alone is shown as a control for signal bleed from the 488 channel. (F) Cells expressing C11orf49-HA-FLAG were fixed and stained with C11orf49 and ᵞ-tubulin endogenous antibodies. A single plane of a 2 µm z-stack is shown; scale bar represents 20 µm.

**Table S1: BioPlex 3.0 Network and Analysis, Related to** **Figure 1|** (A) List of baits targeted for AP-MS in 293T cells (B) List of interactions observed in 293T cells (C) – (I) Results from enrichment analysis of BioPlex 3.0 network targeting GO Biological Process, Molecular Function, and Cellular Components, Reactome pathways, PFAM and InterPro domains, and DisGeNET diseases. (J) Results from prediction of protein subcellular localizations based on BioPlex 3.0 network structure. (K) Summary of communities identified in the BioPlex 3.0 network via MCL clustering (L) Summary of proteins assigned to each community via MCL clustering. (M) List of PFAM domain-domain associations identified in the 293T network.

**Table S2: BioPlex HCT116 Network, AP-MS Replicates, and Quantitative Proteomic Comparison of Cell Lines, Related to Figures 2, S1, and S2**. (A) Data from RTS-MS3 TMT analysis of 293T and HCT116 proteomes in biological quintuplicate. (B) List of all baits targeted for AP-MS in HCT116 cells. (C) List of all interactions observed in HCT116 cells via AP-MS. (D) Merged list of interactions observed in either 293T cells, HCT116 cells, or both. (E) Summary of 293T replicate AP-MS experiments. (F) Summary of interactions observed in 293T replicate experiments. (G) Summary of HCT116 replicate AP-MS experiments. (H) Summary of interactions observed in HCT116 replicate experiments. (I) Summary of 293T-HCT116 replicate AP-MS experiments. (J) Summary of interactions observed in 293T-HCT116 replicate experiments.

**Table S3: Interaction Overlap among Functional Subnetworks, Related to** Figures 3 and S5. (A) Summary of edge sharing between 293T and HCT116 cells within subnetworks defined by specific DisGeNET disease associations. Only edges detectable in both cell lines are considered. (B) Summary of edge sharing between 293T and HCT116 cells within subnetworks defined by specific GO Biological Processes. Only edges detectable in both cell lines are considered. (C) Summary of edge sharing between 293T and HCT116 cells within subnetworks defined by specific GO Molecular Functions. Only edges detectable in both cell lines are considered. (D) Summary of edge sharing between 293T and HCT116 cells within subnetworks defined by specific GO Cellular Components. Only edges detectable in both cell lines are considered. (E) Summary of edge sharing between 293T and HCT116 cells within subnetworks defined by specific Reactome pathways. Only edges detectable in both cell lines are considered. (F) Summary of edge sharing between 293T and HCT116 cells within subnetworks defined by specific CORUM complexes. Only edges detectable in both cell lines are considered.

**Table S4: Discovery and Cell-line Specificity of BioPlex Communities, Related to** Figure 4. (A) Summary of Communities identified in the combined BioPlex 293T + HCT116 network. (B) List of Entrez Gene ID’s and Symbols for proteins found in each community. (C) List of community pairs whose members interact with each other unusually often, along with p-values reflecting the strength of the association between communities. (D) Summary of the fraction of interactions within each community that are shared between cell lines. Only edges detectable in both cell lines are considered. (E) Summary of the fraction of interactions linking each community pair that are shared between cell lines. Only edges detectable in both cell lines are considered.

**Table S5: PFAM Domain Associations and Functional Associations in the Combined BioPlex Network, Related to** Figures 5 and S6. (A) A summary of all PFAM domain associations in the combined 293T/HCT116 BioPlex network. (B) For each domain association, the fraction of edges that are specific to 293T or HCT116 cells or detected in both. Only edges detectable in both cell lines are considered. (C) For baits targeted in both 293T and HCT116 cells, a list of enriched terms (GO, PFAM, InterPro) and the fraction of matching proteins that are unique to 293T or HCT116 cells or detected in both.

**TABLE S6: Identifying Accessory Proteins for Known Protein Complexes, Related to** Figures 6 and S7. (A) Summary of CORUM complexes considered for scoring. We required 4+ members overall, 4+ members detected in both cell lines, and 2+ baits. (B) Summary of proteins scored for association with each CORUM complex. Known complex members were scored via leave-one-out cross-validation. (C) Summary of proteins scored for association with the Tubulin Polyglutamylase Complex. Known members were scored via leave-one-out cross-validation.

**TABLE S7: BioPlex and Achilles: Linking Physical and Functional Associations for Biological Discovery, Related to** Figure 7. (A) List of fitness correlations calculated for those edges in the combined BioPlex network where Achilles data were available for both interacting proteins.

